# Long noncoding RNA *VENTHEART* is required for cardiomyocyte specification and function

**DOI:** 10.1101/2021.02.01.429136

**Authors:** Albert Dashi, Wilson L.W. Tan, Chukwuemeka George Anene-Nzelu, Bangfen Pan, Autio Matias Ilmari, Zenia Tiang, Robin J.G. Hartman, Justus Stenzig, Heming Wei, Chen Gao Bin, Matthew Andrew Ackers-Johnson, Bing Lim, Anna Walentinsson, Vidhya Vardharajan Iyer, Malin K.B. Jonsson, Roger S. Foo

## Abstract

**Rationale:** Long noncoding RNAs (lncRNAs) control cardiac gene expression during heart development and disease. It is accordingly plausible for the same lncRNA to regulate both cardiac development, as well as play a role in adult heart disease progression. lncRNA regulators of early cardiomyocyte (CM) lineage commitment have been identified and characterised, however those controlling later CM specification remain unknown.

**Objectives:** In this study we identified a novel lncRNA required for CM specification, maturation and function, and also discovered its suggested relevance to heart disease.

**Methods and Results:** We performed single cell RNA-seq on human embryonic stem cell derived cardiomyocytes at 2, 6 and 12 weeks of differentiation. Weighted correlation network analysis (WGCNA) identified core gene modules, including lncRNAs highly abundant and uniquely expressed in the human heart. A lncRNA (we call *VENTHEART*, *VHRT*) co-expressed with cardiac maturation and ventricular-specific genes *MYL2* and *MYH7*, as well as in adult human ventricular tissue. CRISPR-mediated excision of *VHRT* led to impaired CM sarcomere formation, and loss of the CM specification gene program. *VHRT* knockdown (KD) in hESC-CMs confirmed its regulatory role for key cardiac contraction, calcium hemostasis and heart development genes, including *MYH6* and *RYR2*. Functional evaluation after *VHRT* KD using impedance-based technology and action potential recordings, proved reduced contraction amplitude and loss of the ventricular-like action potential in CM, respectively. Through an integrative analysis of genome-wide association studies (GWAS), expression quantitative trait locus (eQTL) and gene co-expression network, we found *VHRT* to be co-regulated with core cardiac contractile genes, and the likely source of a heart failure genetic association signal overlapping the *VHRT* gene locus. Finally, *VHRT* KD and human failing heart transcriptome comparison validates the consistent downregulation again of cardiac contractile and calcium regulatory genes (*P*<0.05).

**Conclusion:** We conclude that *VHRT* lncRNA is required for proper CM specification and function. Furthermore, reduced *VHRT* may contribute to the development or progression of human heart disease.

## INTRODUCTION

Cardiac development and heart disease progression are precisely coordinated at the transcriptome level where gene expression programs are jointly controlled through signaling pathways and epigenetic regulators. Human embryonic stem cell-derived cardiomyocytes (hESC-CMs) are a useful tool to study cardiac commitment through their defined stages of cardiac differentiation^1, 2^. This *in vitro* system can model cardiac diseases including various forms of long QT syndrome^3^, arrhythmogenic right ventricular dysplasia^4^, hypertrophic cardiomyopathy^5, 6^ and ischemic heart conditions^7^. Furthermore, hESC-CM are routinely used to test novel drugs or therapeutic options. As examples, RNA interference (RNAi) rescued the disease phenotype of hESC-CMs carrying a mutation causing long QT syndrome^8^, or carrying a mutation in phospholamban that results in dilated and arrhythmogenic cardiomyopathy^9^.

As an *in vitro* cell model, many studies have investigated the mechanisms underlying early cardiac commitment from hESC through to mesoderm and cardiac progenitor stages, and the signaling pathways responsible for these transitions are relatively well-defined^10, 11^. However, the later stages of CM specification and maturation are less well-studied. Other than transcription factors, chromatin modifiers and microRNAs that are well-known regulators of transcription, long noncoding RNAs (lncRNAs) represent an additional layer of regulation that coordinate cardiac gene programs. LncRNAs are >200 nucleotide long transcripts with limited protein coding potential, and often display species, tissue- and disease-specific expression patterns^12^. They are diverse in their cellular expression patterns, subcellular localization, evolutionary conservation, and mechanisms of action. Thousands of human cardiac lncRNAs have been catalogued^13^, but most remain to be functionally characterized. Previous examples for cardiac development are *Braveheart (Bvht*), necessary for the progression of nascent mesoderm towards a cardiac fate through two different mechanisms of action^14, 15^, and *Fendrr*, specifically expressed in the nascent lateral plate mesoderm and essential for proper development of the heart and body wall^16^. More recently, another lncRNA was found to regulate murine mesodermal specification by recruiting transcription factors eomes, trithorax group (TrxG) subunit WDR5, and histone acetyltransferase GCN5 to the enhancer region of *Mesp1* gene, directly activating its expression^17^. These studies were implemented using global transcriptome analysis focused on early cardiac lineage commitment. Transcriptome profiling of the developing mammalian heart has also revealed lncRNAs uniquely expressed in certain sets of cells and activated upon specific stimuli^18–20^. As with early differentiation, distinct lncRNA expression patterns in specific stages and cell states of CM specification and maturation also underlie their essential biological function in heart disease.

Here, we have differentiated hESC to CM, and performed single cell RNA-seq analysis for co-expression and transcriptional changes that occurred in hESC-CMs over time in culture. Hypothesizing that key co-expressed lncRNAs could be drivers of CM specification, maturation and function, we identified *VENTHEART* (*VHRT*) that was co-regulated with core cardiac development and maturation genes and highly expressed in ventricular CMs. Additionally, an independent integrative analysis of genome-wide association studies (GWAS), expression quantitative trait locus (eQTL) and human heart tissue transcriptomes also pointed to the role of *VHRT* in cardiac function and heart failure disease association.

## RESULTS

### Single cell RNA-seq of maturing hESC-CM identifies cardiac specific lncRNAs

A human ESC line with a *MYH6*-GFP reporter was generated and differentiated into CMs, using the protocol published by Lian *et al*^21^ with minor modifications **(Figure 1A, Supplementary Figure 1A-D)**. To map the single cell transcriptome of maturing hESC-CM, we captured cells from week 2 (W02), week 6 (W06) and week 12 (W12) using the Fluidigm microfluidic C1 system. A total of 173 GFP+ (CMs) and 35 GFP-(non-CMs) cells from two biological repeats were sequenced. To keep only the most optimum dataset, 21 samples were excluded from analysis due to either inadequate library size (<1 M reads) or a low number of detected transcripts (<2,000 genes expressed) **(Supplementary Tables 1 and 2)**. This resulted in a total of 36 CMs at W2, 60 CMs at W6 and 56 CMs at W12. To assess the validity and quality of our data we undertook several further technical QC procedures **(Supplementary Figure 2A-F, Supplementary Table 1).** Only CMs (GFP+) were used for subsequent analysis for the remainder of this project. To confirm increased hESC-CM maturation, we firstly confirmed the consistent and timely upregulation of selected maturation genes (such as *TRDN*, *MYH7*, *TNNI3* and *MYL2*) **(Supplementary Figure 3A)**, as well as calcium handling genes (*RYR2*, *ASPH*, *PLN* and *ATP2A2*). Notably, genes for cellular coupling gap junctions (*GJC1*), ion channel proteins (*KCNJ2*, *KCNA5*, *CACNA1G* and *HCN4*) and sarcomere assembly (*TCAP*) were also increased.

**Figure 1.**
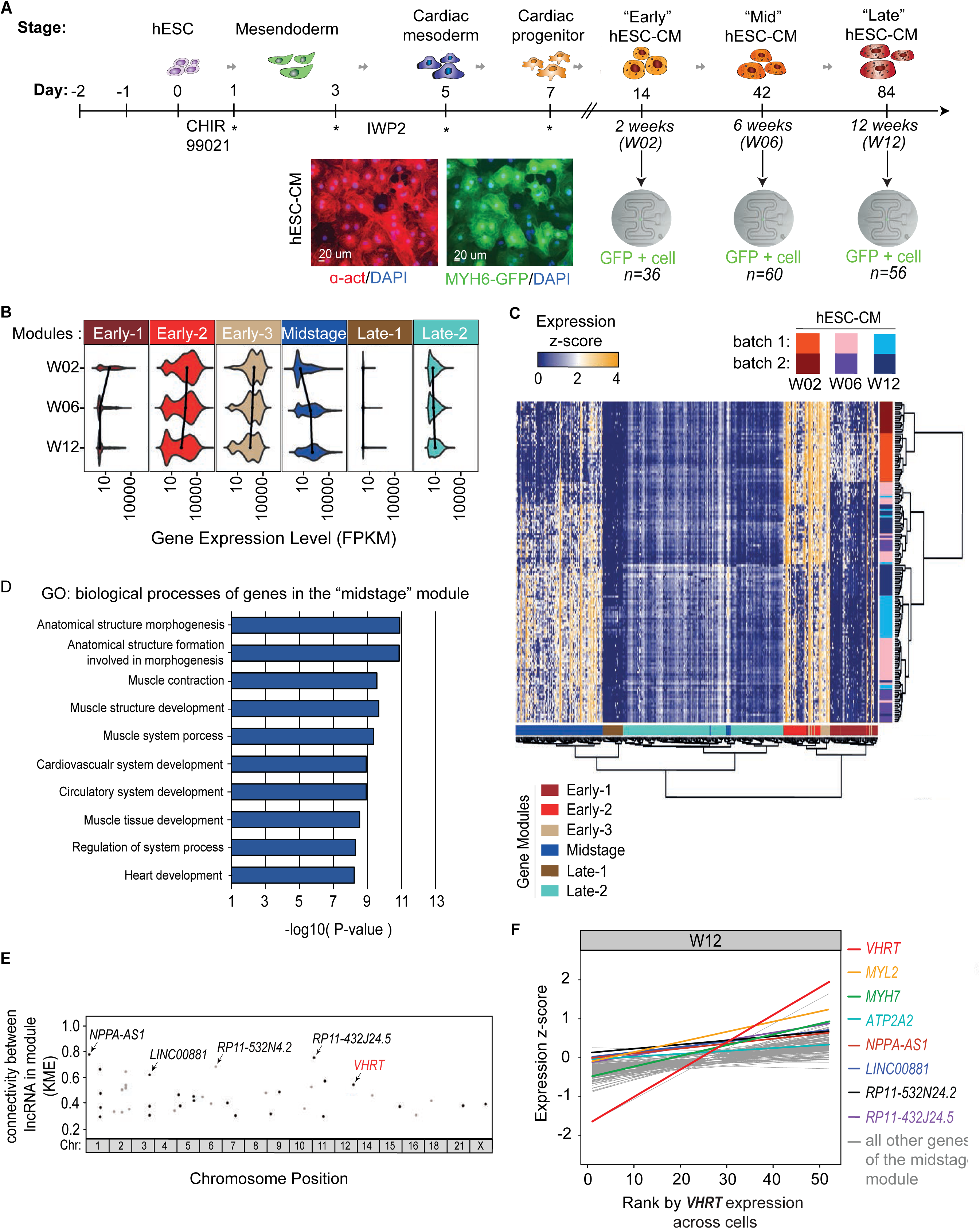
Single cells RNA-seq analysis identifies cardiac enriched lncRNAs in week 2-12 hESC-CM. **(A),** A schematic of the experimental plan. Differentiation of hESC to CM was performed over 7 days under insulin-free conditions and with the addition of small molecule CHIR99012 at day 0 and IWP2 at day 3. Additional media change was performed on days as denoted (*). 36 cells were harvested on week 2 (W02), 60 cells on week 6 (W06) and 56 cells on week 12 (W12). Immunofluorescence staining for cardiac alpha actinin (ACTN2, red) mark sarcomeres, and green represents *MYH6*-GFP reporter signal. **(B),** Violin plots showing the overall levels of gene expression within each gene module. Each of the six modules (early-1, early-2, early-3, midstage, late-1, late-2) shows either overall increase or decrease gene expression over time. Black lines track the median expression values across the time points. **(C),** Heatmap showing the expression of all genes in the six modules with genes in columns. Cells from each time point and replicates are in rows. **(D),** Histogram showing top 10 biological processes of genes in the “midstage” blue module. **(E),** Plot showing the connectivity (KME values) of lncRNAs associated with the other protein coding genes in the “midstage” module. Highly connected lncRNAs with KME values closest to 1 represent those most highly co-regulated with other genes in the module. Five highly co-regulated lncRNAs that are highly expressed in muscle heart are indicated. *RP1-46F2.2* (we call *VENTHEART, VHRT*; on chromosome 12) is annotated in red. **(F),** Plot showing the expression of all “midstage” module genes (expression z-scores) in W12 hES-CMs ranked from 1 to 56 according to *VHRT* expression (red line). *VENTHEART* is correlated with the expression with core cardiac genes *MYL2*, *MYH7*, *ATP2A2,* and other genes of the “midstage” module. Lines are fitted linear for z-score values.

**Table 1:** CardioExocyte profile from CM after VHRT siRNA knockdown. Contractility profile was measured using Cardioexcyte 96 instrument (Nanion). Base impendance, beat rate, amplitude, pulse width, upstroke velocity and down velocity are shown. *ATP2A2* siRNA was used as positive control and non-targeted siRNA (NT) as well as un-treated (UT) as negative controls. The data was analyzed using the data control software from Nanion. One-way ANOVA, multiple comparisons, compared to control siRNA. Data are shown as mean ± s.e.m. * *p*<0.05, \*\**p*<0.0, \*\*\**p*<0.001 and \*\*\*\**p*<0.0001.

|  | Base Impedance<br>[ohm] | Beat rate<br>[bpm] | Amplitude<br>[ohm] | Pulse width 50 %<br>[ms] | Upstroke velocity<br>[ohm/s] | Downstroke velocity<br>[ohm/s] |
| --- | --- | --- | --- | --- | --- | --- |
| UT | 577±18** | 60±0 | 7.1±0.4** | 162±10 | 223±29*** | -56±5 |
| NT | 523±39 | 60±0 | 6.0±0.6 | 164±14 | 176±22 | -50±7 |
| *VHRT* siRNA | 403±8**** | 60±0 | 4.4±0.2**** | 176±8 | 108±3**** | -36±2*** |
| *ATP2A2* siRNA | 405±26**** | 60±0 | 3.3±0.5**** | 260±10**** | 63±9**** | -17±2**** |

In order to explore for unsupervised co-regulated gene expression, we subjected the data to weighted correlation network analysis (WGCNA), and constructed gene-network modules that characterized expression changes across the time-course **(Supplementary Figure 4A, Supplementary Table 3)**. Six modules differed significantly between the time points **(Figure 1B, Supplementary Figure 4B and Supplementary Table 4)**. The “early-1” gene module consisted of genes highly expressed in W2 cells, and significantly downregulated in W6 and W12. Gene ontology (GO) functions for this “early-1” module was related to general tissue development and heart morphogenesis. The second module, “early-2”, was also highly expressed at W2, and decreased at W6 and W12. The “early-2” module included pathways related to protein transport and localization. “Early-3” was another module that showed decreased expression over the time-course and contained GO pathways linked to cell cycle control. A “midstage” module showed the distinct opposite pattern of increasing expression from W02 to W06, and a further increase at W12. Genes in the “midstage” module were related to cardiac muscle development, maturation and cardiac functions, reflecting hESC-CMs strengthening their cardiac identity after W2 and mature and specify over time. Two further “late” modules showed a statistically significant but less marked difference during the time-course. “Late-1” included genes with general functions such as tau protein binding, “late-2” were those involved in general metabolic processes. Genes in all six modules reliably segregated cells into the three time points (**Figure 1C**), giving a comprehensive single cell transcriptional landscape of maturing hESC-CM over this time course.

Next, we curated for lncRNAs that correlated with transcriptional changes, and paid specific attention to the “midstage” blue module with gene pathways involving cardiac muscle development and maturation processes, including sarcoplasmic reticulum ion transport and actin mediated cell contraction (**Figure 1D**). In this module we identified a set of lncRNAs highly co-regulated with other important cardiac specific genes (**Figure 1E**). Among them, three (*RP11-532N4.2*, *LINC00881* and *RP11-432J24.5*) are highly expressed in human hearts, and one (*RP1-46F2.2,* hg19 or *LINC01405*, hg38) is limited to only cardiac ventricle and skeletal muscle, as shown from the publicly available human tissue specific expression dataset (GTEx)^22, 23^ **(Supplementary Figure 4C)**. Specifically, *LINC01405* we now call *VENTHEART* (*VHRT*), followed the tissue expression pattern of sarcomere assembly and calcium regulatory genes, including myosin light chain 2 (*MYL2*), myosin heavy chain beta (*MYH7*) and sarco/endoplasmic reticulum Ca2+-ATPase (*ATP2A2*), and showed the steepest increase over the time-course compared to the other lncRNAs (**Figure 1F**). Human ESC-CMs in our 12-week time-course were largely negative for markers of the secondary heart field (*ISL1* and *KDR*), and instead positive for primary heart field markers, *TBX5* and *CORIN*^18^, and were MLC2v+ (encoded by *MYL2*) for ventricular-like CMs **(Supplementary Figure 3B-C)**. Indeed ranking all W12 hESC-CM samples from low to high *VHRT* expression, confirmed that the expression of *MYL2*, *MYH7*, *ATP2A2* and other genes of the same module (“midstage”) was co-linear with *VHRT* (**Figure 1F**), suggesting again that there are shared regulatory properties between genes in this module.

### *VENTHEART* is highly enriched in ventricular-like CMs

*VHRT* is highly conserved only among primates^24^ **(Supplementary Figure 5À)**, and the *VHRT* locus is annotated to comprise of 6 isoforms (**Figure 2A, Supplementary Table 5)**. To confirm differential isoform expression, we performed primer specific RT-qPCR and identified isoform 6 as the most highly expressed in hESC-CMs (**Figure 2B**). Isoform 6 *VHRT* was abundant in the cytoplasmic compartment, whereas the other less expressed isoforms showed an unremarkable distribution across nuclear and cytoplasmic compartments (**Figure 2C, Supplementary Figure 5B)**. For cell and tissue expression specificity, we found *VHRT* to be present in hESC-CM, but absent in human cardiac fibroblasts (hCF), smooth muscle cells (hCASMC), human monocytic cell line (hTHP-1) and human coronary artery endothelial cells (hCAEC) (**Figure 2D**). Furthermore, human heart tissue expression analysis confirmed that *VHRT* is absent or lowly expressed in atria, compared to ventricular tissue **(Supplementary Figure 5C)**. This recapitulated data from GTEx portal, further underlining the specific *VHRT* expression pattern in the heart. The Coding Potential Calculator (CPC)^25^ algorithm suggested that among all isoforms, isoform 6 may possess protein coding potential with a putative 76 aa long peptide^26^**(Supplementary Figure 6A-B)**. We proceeded to test for the putative peptide empirically by FLAG-tag at either C-terminus or N-terminus of *VHRT*-6 sORF (without introns), cloning it into a vector for cell transfection. No bands were detected on Western blotting despite relevant positive controls **(Supplementary Figure 6C)**. This may reflect the possibility that the transcript is not translated into a stable or functional peptide *in vitro*, and *VHRT* may function instead as a *bona fide* non-coding RNA.

**Figure 2.**
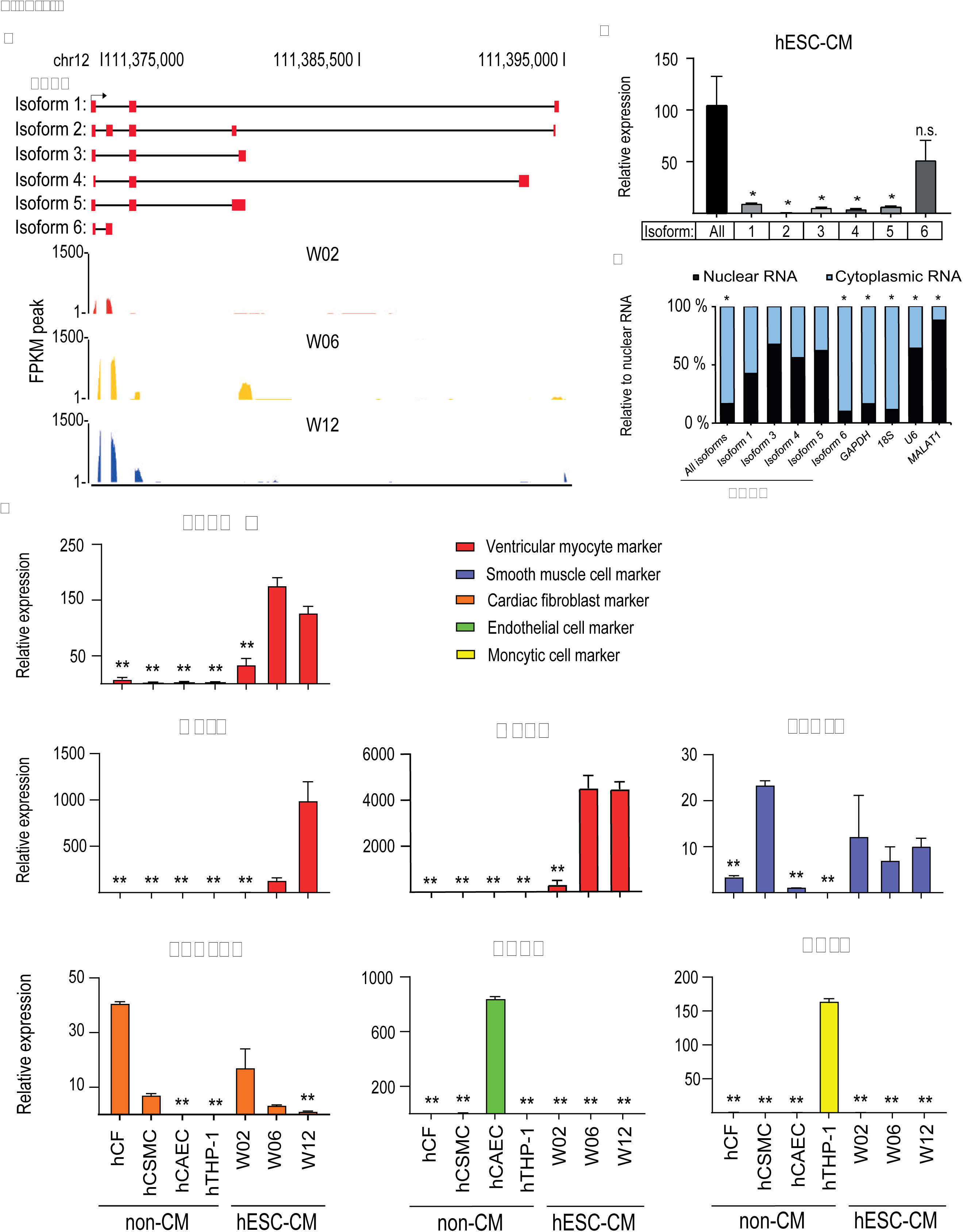
Characterisation of *VENTHEART* expression in human hES-CM. **(A),** Schematic representing the 6 *VHRT* isoforms annotated on NONCODE and ENSEMBL. Top panel, red boxes represent exons. Exon 1 is shared by all isoforms. Lower panel shows normalized FPKM peaks from single cell RNA-seq for W02 (red), W06 (yellow) and W12 (blue), reflecting expression abundance of *VHRT* at each time point. **(B),** Relative quantification of specific *VHRT* isoforms expression. Data are represented as mean ± s.e.m., n=3; * *p*-value < 0.05 are represented; n.s.: not significant. Student’s t-test. **(C),** Graphs showing nuclear and cytoplasmic fractions. *GAPDH* and *18s* used as positive controls for cytoplasmic compartment, and nuclear compartment with *U6* and *MALAT1*. Expression are shown as percentage relative to nuclear fraction. n=3 biological replicates. * represents p ≤ 0.05. Student’s paired t-test with a two-tailed distribution. **(D),** *VHRT* expression is present only in CM, but absent in non-CM. Specific markers were used for the different cell types: *PDGFRA* for cardiac fibroblast (hCF), *TAGLN* for smooth muscle cells (hCASMC), *MYL2* and *MYH7* for hESC-CM and *CDH5* for endothelial cells (hCAEC). Data are represented as mean ± s.e.m., n=3. ** *p*-value < 0.01. Student’s t-test.

### *VHRT* deletion disrupts CM specification

To test the hypothesis that *VHRT* regulates CM differentiation or specification, we performed *VHRT* CRISPR/Cas9-directed knockout (KO) at hESC stage using a dual sgRNA system. To control for off-target effects from CRISPR-targeting, we designed 2 unique sgRNA pairs that targeted different locations of *VHRT (***Figure 3A-B**). Two independent homozygous knockout hESC lines were obtained (clone 1: *VHRT* KO #1 and clone 16: *VHRT* KO #2). Untargeted cells were used as wild type control (clone 9: WT) **(Supplementary Figure 8A-B).** Independent tri-lineage differentiation proved that ectodermal, mesodermal and endodermal lineages were unaffected in both *VHRT* KO lines **(Supplementary Figure 9A)**. We then differentiated WT and KO lines to CM in parallel. Apart from minimum variability between differentiations batches, both KO lines entered the initial stages of differentiation efficiently **(Supplementary Figure 9B)**. Key genes of the cardiac progenitor stage, such as *NKX2-5,* were unchanged. The *MYH6*-GFP+ signal however was significantly reduced in KO at day 7-8, compared to WT **(Supplementary Figure 10A)**, reflecting the likely importance of *VHRT* for the transition from cardiac progenitor to early CM. Remarkably, upon continued culture, *VHRT* KO cells showed reduced CM specification and maturation genes (**Figure 3C**), and an inability to adequately develop sarcomere structure and organization from W6 to W12. Hence, while WT hESC-CM strengthened their characteristic cellular organization from W2 to W12, the lack of proper sarcomere assembly in *VHRT* KO #1 and *VHRT* KO #2 may reflect a failure to activate CM gene programs **(Figure 3D, Supplementary Figure 10B-C).**

**Figure 3.**
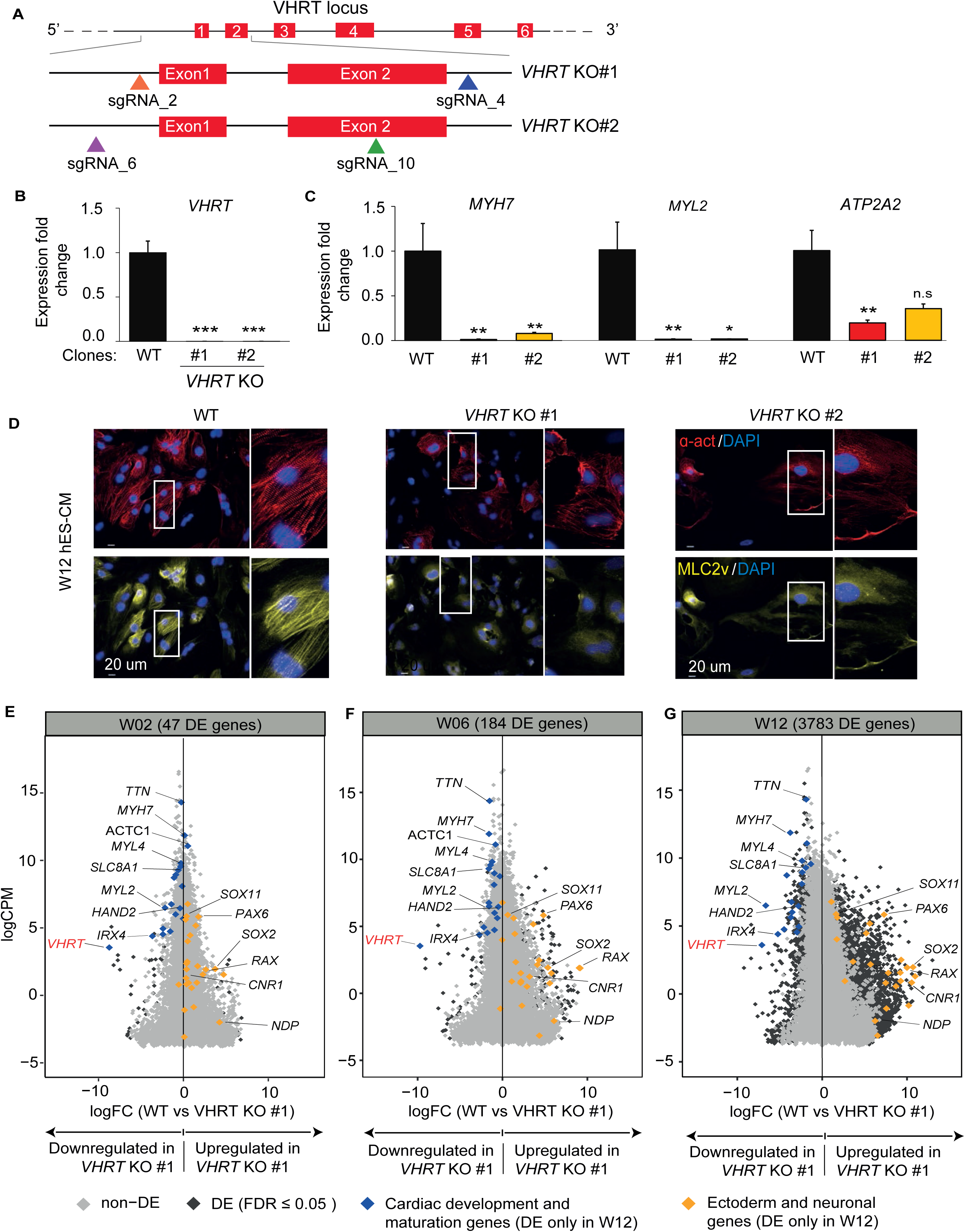
CRISPR-mediated *VENTHEART* deletion results in impaired CM specification. **(A),** Schematic showing CRISPR/eSpCas-9-mediated *VHRT* knockout (KO). First hESC KO line (*VHRT* KO #1) was generating using sgRNA_2 and sgRNA_4, and the second (*VHRT* KO #2) using sgRNA_6 and sgRNA_10. **(B),** RT-qPCR validating loss of *VHRT* expression in *VHRT* KO #1 and #2. Expression is represented as fold change relative to wild type (WT). **(C),** Significant downregulated expression of cardiac genes *MYH7*, *MYL2* and *ATP2A2* in W12 *VHRT* KO cells. **(D),** Representative immunostainig showing sarcomere disorganization for *VHRT* KO #1 and #2, compared to WT in W12 hES-CM. **(E-G),** Distribution of gene expression log fold change (logFC), comparing between GFP+ cells in WT and *VHRT* KO #1 for W02 **(E)**, W06 **(F)** and W12 **(G).** Differentially expressed (DE) genes at each time point are highlighted in dark-grey. Non-DE genes in light-grey. Highlighted in blue are cardiac development and maturation genes from the “midstage” module in Figure 1D, significantly downregulated in W12. Orange are ectoderm and neuronal genes, significantly upregulated in W12. Data are represented as expression fold change ± s.e.m., n=3. ** *p*-value < 0.01, \*\*\**p*-value < 0.001, n.s.: not significant. Student’s paired t-test with a two-tailed distribution.

### *VHRT* deletion deregulates the transcriptome of maturing hESC-CM

A small subset of *VHRT* KO cells (15% of *VHRT* KO #1, and 20% of *VHRT* KO #2) turned on the *MYH6*-GFP reporter during the differentiation protocol **(Supplementary Figure 11A)**. This raised the question whether some *VHRT* KO cells developed general CM gene expression. We therefore isolated GFP+ cells from WT and *VHRT* KO #1 at W2, W6 and W12 **(Supplementary Figure 11B)**, and performed RNA-seq to compare global gene expression. *VHRT* deletion resulted in a global increase in differentially expressed genes over time in culture (**Figure 3E-G**). Across W2, W6 and W12, cardiac contractile and calcium regulatory genes including *MYH7, MYL2, SLC8A1* and *IRX4* and failed to upregulate in *VHRT* KO, compared with WT **(Figure 3C-E, Supplementary Table 6)**. In contrast, ectodermal and neuronal function genes, such as *SOX2, PAX6* and *RAX*, were significantly upregulated, especially in W12 *VHRT* KO cells (**Figure 3G, Supplementary Table 6)**. Furthermore, genes involved in metabolic processes were persistently downregulated from W2 to W12, reflecting the lack of a maturation metabolic switch in *VHRT* KO **(Supplementary Table 6)**.

### *VHRT* knockdown downregulates sarcomere assembly and calcium handling genes, and reduces CM contractility

Next, we sought to assess whether *VHRT* is also implicated in the regulation and maintenance of CM function in hES-CM. We therefore performed *VHRT* knockdown (KD) using 2 independent GapmeRs, specifically targeting *VHRT* isoform 6, since it is the most expressed isoform in hES-CMs (**Figure 4A**). This was carried out in W6 hESC-CM, where *VHRT* gene expression was elevated compared to preceding stages of differentiation. 75% *VHRT* KD was achieved with GapmeR#1, and 72.4% with GapmeR#2, respectively. (**Figure 4B**).

**Figure 4.**
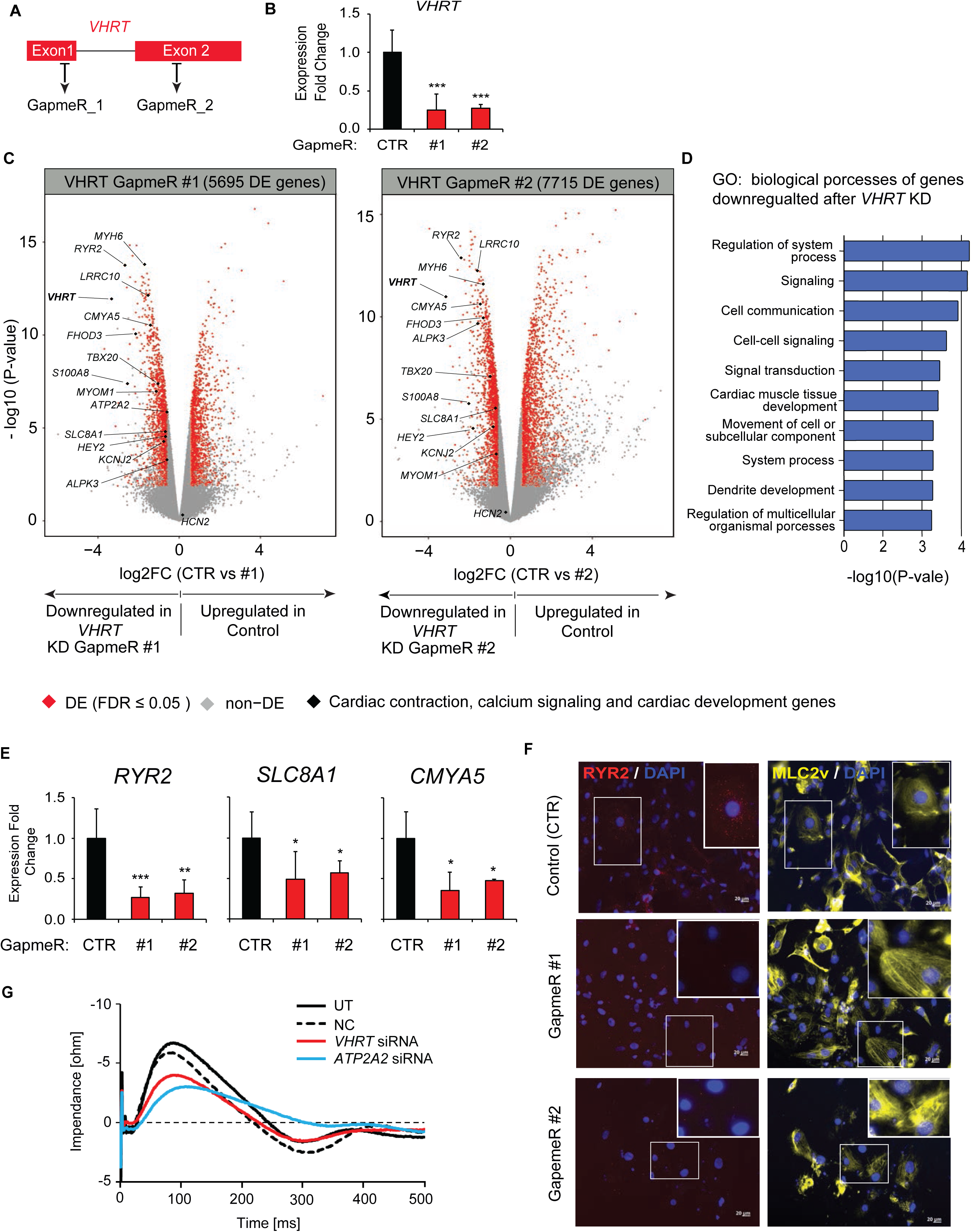
*VENTHEART* knockdown decreases core cardiac gene expression and disrupts CM function. **(A),** Schematic showing targeting GapmeRs. **(B),** RT-qPCR validation for *VHRT* knockdown (KD) in W06 hESC-CM, transfected with GapmeR#1 (n=5), GapmeR#2 (n=3) or GapmeR non-targeting control (CTR, n=5), and harvested for analysis after 4 days. **(C),** Volcano plot showing DE genes for *VHRT* GapmR#1 (left panel) and *VHRT* GapmeR#2 (right panel). Highlighted in red are significant DE genes. Non-DE genes are marked in grey. In black are annotated cardiac contraction, calcium signaling and cardiac development genes significantly downregulated by *VHRT* KD. **(D),** Histogram showing top biological process downregulated after *VHRT* GapmeR-mediated KD. **(E),** Significant downregulation of calcium handling genes *RYR2*, *SLC8A1* and validated by RT-qPCR. Expression are shown as relative expression to control (CTR). n=3 biological replicates. *p ≤ 0.05, ** p ≤ 0.01 and ***p ≤ 0.001. Student’s paired t-test with a two-tailed distribution. **(F),** Immunostainig showing reduced ryanodine receptor 2 (RYR2) protein expression in *VHRT* GapmeR#1 KD and #2 KD, compared to CTR, in W06 hES-CM. **G)**, Impedance-based contractility profile analysed using the CardioExcyte (Nanion). Reduction in relaxation velocity, upstroke velocity, amplitude and prolonged pulse width were observed for *VHRT* siRNA, similar to *ATP2A2* siRNA as positive control.

*VHRT* KD CM showed significant differentially expressed (DE) genes belonging to biological processes controlling cardiac electrophysiology, cardiac structure and heart development (**Figure 4C-D**). There was significant downregulation of structural and sarcomere assembly genes including *MYH6*, *FHOD3* and *MYOM1,* and known markers of heart tissue morphogenesis *TBX20*, *HEY2* and *ALPK3*. Remarkably, a strong repression was observed for calcium regulatory genes including *RYR2, SLC8A1* and *CMYA5* (**Figure 4E-F**). Importantly, these findings were specific because the same significant changes were not observed for other CM marker genes (**Figure 4B-D**, non-DE genes). To assess the function on contractility and CM electrophysiology, we also performed an independent *VHRT* siRNA KD, followed by impedance-based assays. *VHRT* KD significantly affected contractility performance, with decreased relaxation velocity, decreased upstroke velocity, decreased amplitude and a prolonged pulse width, compared to non-targeted siRNA (NT) and untreated (UT) controls (**Figure 4G, Supplementary Figure 11A)**. *VHRT* KD induced similar changes in contractility as KD of *ATP2A2,* encoding for SERCA2a, consistent with altered calcium handling in *VHRT*-deficient CM. **(Figure 4G, Table 1).** Patch clamp experiments showed that *VHRT*-KD cells had significantly more positive maximum depolarization potential (MDP), compared to control (−54.2±6.5 vs −61.2±7.1 mV, n=28, *p*≤0.01). *VHRT*-KD cells also displayed a more triangular action potential (AP) waveform, as assessed by either the APD90/APD50 ratio (1.34±0.12 vs 1.23±0.14, p≤0.05) or the APD90-APD20 difference (256±81 vs 166±50 ms, *p*≤0.001) **(Supplementary Figure 11B).** Representative AP from spontaneously beating cells and cells paced at 1Hz are shown in **Supplementary Figure 11C**. Taken together, these data confirmed that *VHRT* KD significantly compromised CM function.

### Integrated GWAS and eQTL analysis implicates *VHRT* in human heart disease

Related to the unique expression profile of *VHRT* for differentiating CMs above, and the well-known hallmark of “fetal gene reprogramming” in the heart disease stress-gene response, we also set out to investigate a potential role for *VHRT* in human heart disease, via analysis of publicly available genome-wide association study (GWAS) data sets^27–31^. Indeed, at least 2 independent GWAS signals were located in or near the *VHRT* locus at chromosome 12q24.11. Significant signals (*p*-value<5e-8) for coronary artery disease (CAD) and myocardial infarction (MI) overlapped with the genes *SH2B3* and *ATXN2* downstream of *VHRT*, but these were unlikely related to *VHRT* as they did not associate with cardiac *VHRT* expression based on cis-eQTL data from human left ventricular (LV) samples.

Instead, a tightly linked cluster of SNPs (r2>0.8) overlapping with the *VHRT* gene and significantly associated with *VHRT* expression in heart LV, showed suggestive association with non-ischemic heart failure (HF) (*p*-value=5e-5) (**Figure 5A, B**), indicating that the HF GWAS signal is most likely mediated by altered *VHRT* cardiac expression. SNPs associated with reduced cardiac *VHRT* expression were associated with increased risk for HF (OR=1.12-1.15) **(Supplementary Figure 12A-B)**. Importantly, none of the SNPs showed association with *MYL2* cardiac expression, suggesting that *VHRT* rather than *MYL2* is the likely mediator of the observed cardiac disease GWAS signal. In line with these findings, cardiac *VHRT* mRNA expression was found to be downregulated in a transcriptomics study of patients with dilated cardiomyopathy, compared to healthy controls (adjusted *p*-value<0.05)^31^ (**Figure 5D**). We also performed co-expression analysis using GeneNetwork v2.0^30^, which predicted pathway and human phenotype associations using 31,499 public human RNA-seq samples. *VHRT* was once again tightly co-regulated with key CM sarcomeric genes, including *MYL2*, *MYL3*, *MYL1*, *MYH7*, *TCAP*, *CSRP3* and *TNNC1,* replicating the empirical evidence from the gene network analysis of single cell transcriptomes above. Furthermore, *VHRT* was predicted to participate in heart disease pathways and phenotypes including hypertrophic and dilated cardiomyopathy (HCM and DCM, respectively), as well as ventricular tachycardia **(Figure 5C, Supplementary Table 8)**.

**Figure 5.**
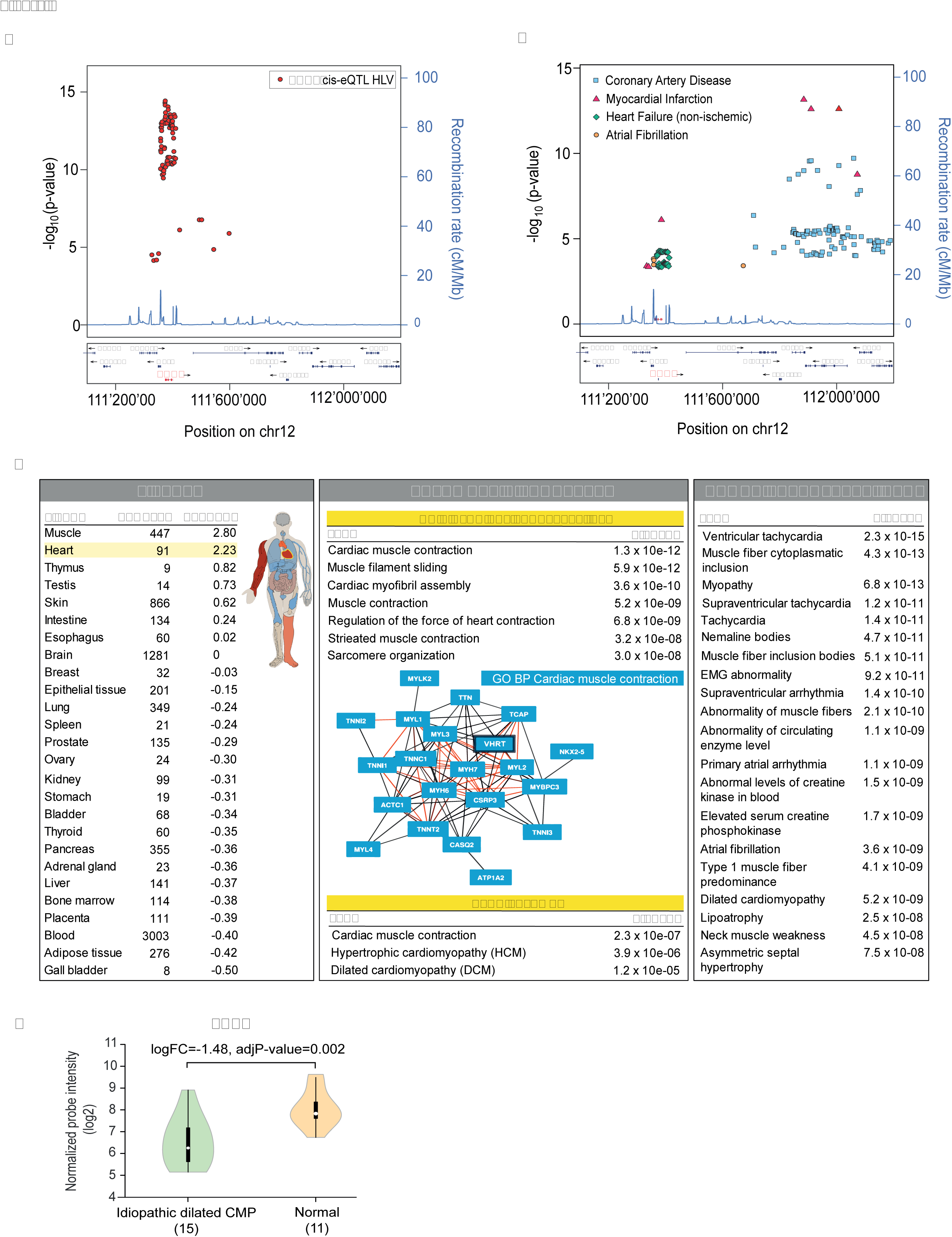
Genetic evidence of a role for *VENTHEART* in heart disease. **(A),** LocusZoom^54^ of 1-MB genomic region harbouring *VHRT* depicting significant *VHRT* cis-*eQTL* SNPs in heart left ventricle (HLV). **(B),** SNPs associated with coronary artery disease (CAD), myocardial infarction (MI), non-ischemic HF and atrial fibrillation (AF) from published GWAS^27–31^. A tightly linked cluster of SNPs (r2>0.8) located at and near the *VHRT* locus shows suggestive association with HF (*P*-value< 5e-5). The same SNPs are significantly associated with *VHRT,* but not *MYL2,* cardiac expression suggesting that *VHRT* rather than *MYL2* is the likely mediator of the observed cardiac disease GWAS signals. **(C),** *VHRT* in cardiac muscle contraction and disease based on co-expression gene network analysis using RNA-seq data from 31,499 human samples^30^. *P*-value < 0.05, Wilcoxon Rank-Sum Test. **(D),** Decreased *VHRT* expression in idiopathic dilated cardiomyopathic failing hearts, compared to healthy controls, based on a re-analysis of microarray data from GEO study GSE1145^31^. Benjamini-Hochberg adjusted *P*-value < 0.05 from univariate analysis with moderated t-statistic using the R limma package, R v3.2 and limma v3.26.0.

### *VHRT* KD transcriptome partially resembled that of gene expression changes in human heart failure

Building on the findings described above, we compared the *VHRT* KD transcriptome of DE genes (N=1,100; 573 down- and 527 upregulated) with the expression profile of the same genes in human failing hearts (Dilated Cardiomyopathy, DCM) (GEO accession: GSE141910). The latter study comprised of RNA-seq from 61 left ventricle explants of idiopathic DCM failing (n=29) and non-failing hearts (NF, n= 32) **(Supplementary Table 9).** Among *VHRT* KD DE genes, 73 (out of 1,100) were also differentially expressed in DCM hearts (**Figure 6A, Supplementary Figure 13**, **Supplementary Table 10)**. Importantly, genes that were downregulated in *VHRT* KD, and recapitulated in DCM (“Down/Down” group) contained those enriched in pathways regulating cardiac muscle contraction, hypertrophic cardiomyopathy (HCM) and dilated cardiomyopathy DCM (p ≤ 0.05), based on pathway and molecular function enrichment analysis using Enricher^32^ (**Figure 6B**). Functional terms related to processes involved in actin filament binding and calcium handling (p ≤ 0.05), including genes such as *MYH6*, *CAMK1D*, *MYO7A* and *CACNB2* **(Figure 6C, Supplementary Figure 13)**, pointing again to the potential mechanisms by which *VHRT* mediates disease progression in HF.

**Figure 6.**
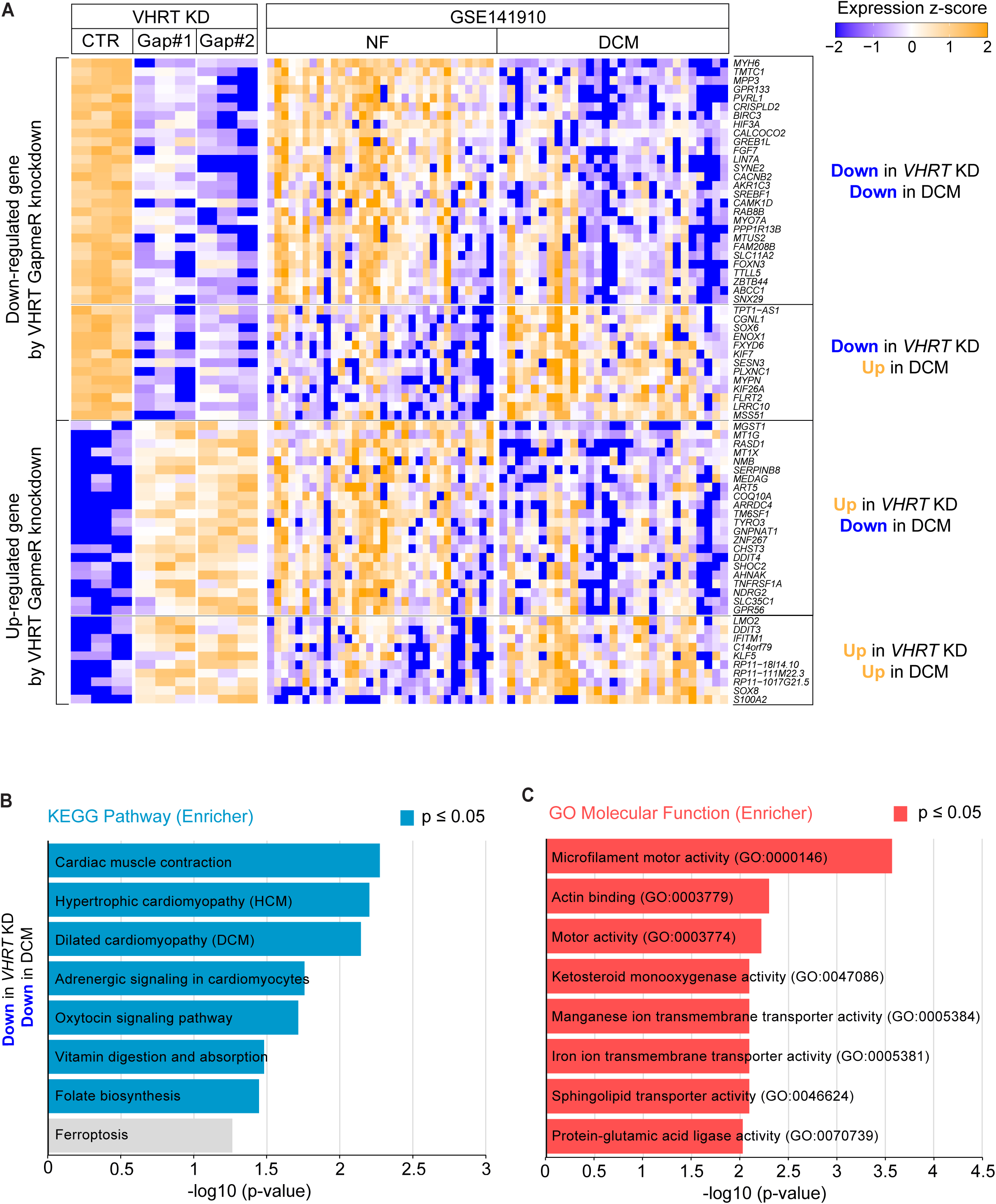
Transcriptome comparison of *VHRT* KD and human failing hearts proposes the gene program mechanism for disease causality. **(A)**, Heatmap comparing *VHRT* KD DE genes with genes that are differentially expressed in a panel of DCM failing hearts (GSE141910) (adjusted *P*-value<0.05). **(B)**, KEGG pathway and (c) Gene Ontology analysis by Enrichr^32^ (log10 (p-value) ≥ 1.3, *p*-value ≤ 0.05) identifies cardiac contractile and calcium regulatory genes consistently downregulated in *VHRT* KD and DCM (“Down/Down”).

## DISCUSSION

Our findings provide the first identification of the long non-coding RNA *VHRT* for CM specification, maturation, and its potential involvement in HF disease pathophysiology. By tracking single-cell resolution transcriptomes of hESC-CMs in prolonged culture, we have compiled an extensive catalogue of hESC-CM expressed genes. Co-expression gene networks, and integrative GWAS and eQTL analyses, highlighted *VHRT* as a novel ventricular-enriched lncRNA, whose expression is limited only to primates.

The overall immaturity of hESC-CMs in culture is well acknowledged. Simple extended time in culture is a useful means to achieve some additional maturity^1, 19, 33, 34^. Our data reveals that key genes, *TNNI3*, *KCNJ2*, *CASQ2*, and *TCAP* are upregulated during CM maturation, appearing at W6 or W12. Between the 2 potassium channel genes: *KCNJ2,* which is ventricular-enriched, and *KCNA5,* which is predominantly atrial, we found that *KCNJ2*+ and *KCNA5*+ hES-CMs are mutually exclusive. Dichotomy for hESC-CMs in this differentiation protocol exists, at least in this respect^35, 36^. We have thus observed that with the 12-week time-course in this differentiation protocol, hESC-CMs come to display largely primary heart field markers (*TBX5*, *HCN4* and *CORIN*), specifying mainly into ventricular-like cells, also corroborated by upregulated ventricular-enriched *MYL2*, *MYH7* and *IRX4*. This is consistent with the ventricular phenotype also described previously for this protocol^21^. Genes in the “early-1” gene module, abundant at W2, and subsequent downregulated in W6 and W12, included those that play a role in general organ development (*BMP2, FGF10, WNT11, TWIST1, SOX11* and *MEF2C*), indicating that even though W2 hESC-CMs are committed to the cardiac lineage, they retain an overall generic transcriptome similar to cells in other organs. Similarly, genes in the “early-2” and “early-3” modules comprised of those for protein transport and localization, and cell cycle regulators (*PSMA6*, *PSMB5*, *PSMA3* and *PSMB4*) that regulate mitotic cell cycle G1/S checkpoint and DNA damage control. The downregulation of these genes coheres with the notion that hESC-CMs decrease their proliferative capacity over the differentiation time-course. In contrast, genes in the “midstage” module are significantly upregulated between W2 and W6, remaining stably abundant thereafter. These are genes related to cardiac development and maturation, including those responsible for cardiac contractile structures (*MYH7, TTN, MYL2, TNNI3, ASPH, MYBPC3*), higher-order myofibrillar organization, cytoskeletal assembly (*MURC132* and *LDB3133*), and calcium handling and transport (*RYR2, ATP2A2, CASQ1, ASPH, ANK2 and SLC8A1*). The “late-1”’ and “late-2” modules contained genes that are only expressed in subsets of W12 cells, where their expression contributed to significant heterogeneity observed at the W12 stage. The “late-2” module comprised of a long list of ribosomal protein pseudogenes associated with general metabolic processes. The significance of the latter will need further investigation.

LncRNAs act as fine switches to modulate and orchestrate multiple aspects of cardiac development. In recent years, many cardiac lncRNAs have been catalogued and functionally characterized, especially for CM differentiation^37–39^. However, less is known for the later stages of CM specification and maturation. Moreover, finding lncRNAs that exclusively regulate either atrial, ventricular or pacemaker cells functions is challenging since different cardiac developmental stages are dictated by different epigenetic mechanisms. The “midstage” module contained specific cardiac protein coding genes known to support CM specification, maturation and function. Therefore, the transcriptional profile of this module accordingly identified CM specific and enriched lncRNAs, among which *VHRT* was highly co-regulated with *MYH7, MYL2, ATP2A2* and other cardiac maturation genes. Being enriched in human ventricular tissue, but not expressed in atrial tissue or other cell types of the heart, *VHRT* may indeed have a unique role in ventricular CM.

CM maturation and specification processes are promoted by pathways and gene programs for metabolism, myofiber structure, electrophysiology and cell cycle^40^. Myosin heavy-chains (MYH6 and MYH7) and light-chains (MYL2) are myofiber proteins important for heart morphogenesis. Pathogenic *MYH7* mutations are causal to hypertrophic cardiomyopathy^41^. Patient-specific iPSC-CMs carrying the *MYH7*-E848G mutation, result in dysregulated myofibril alignment and contractile impairment^42^. In the same way, Wei Zhou et al.^43^ differentiated hiPSC derived CM from patients with *MYL2*-R58Q^44, 45^ mutation and demonstrated that after 60 days in culture, *MYL2*-R58Q-iPSC-CMs also showed sarcomeric disorganization, calcium dysregulation and contractility malfunction. *VHRT* KO, starting from W6, displayed the similar myofilament disassembly and contractility defect, concurrent with the downregulation of cardiac contractile and calcium regulatory genes. Indeed, *VHRT*-KO phenocopies *MYL2*-R58Q and *MYH7*-E848G mutant CMs, implying that the overall deregulated CM gene programme caused by *VHRT*-KO is at least in part, similar to that regulating *MYL2* and *MYH7.* Importantly, pluripotency, mesendoderm and cardiac mesoderm stages were not affected by *VHRT* KO. This was also demonstrated by the finding that general CM genes were normally expressed at W2, whereas with time in culture, ectodermal and neuronal function genes became upregulated instead. However, unlike gene profiles of mutant *MYH7*^46, 47^ and *MYL2* cells^48^, the overall gene profile of W6-12 *VHRT*-KO (and KD) consistently show a downregulation of contractile and calcium regulatory genes, and the lack of a metabolic switch towards “mature” CMs. These results suggest a role for *VHRT* as an epigenetic switch for CM gene programmes during specification and maturation, instead of merely conferring effector functions like those of the sarcomeric components like *MYH7* or *MYL2*.

Other lncRNAs have also been shown to control cardiac gene expression previously. Many do so by interacting with repressive protein complexes and transcription factors in the nucleus, thereby regulating local gene expression through chromatin remodelling. Our characterization experiments however, show *VHRT* to be localized in CM cytoplasm, indicating that *VHRT* may regulate gene programs from the cytoplasmic compartment and not the nucleus, unlike mechanisms for the majority of other lncRNAs. At least from our current assessment, *VHRT* does not appear to encode any detectably stable micropeptide in hES-derived CM. Instead, *VHRT* KD by GapmeR was sufficient to downregulate the expression of cardiac contraction, calcium transport and other ion channel genes. Whole cell patch clamp recordings confirmed that *VHRT* KD cells showed the loss of a ventricular-like action potential. Consistent with functional cytoplasmic *VHRT,* alternative siRNA-mediated *VHRT* KD also verified reduced CM relaxation velocity, upstroke velocity and contractility amplitude. Some lncRNAs are translocated out of the nucleus and shuttled to specific cytoplasmic locations to regulate protein expression or function. Some do so by sponging miRNA^49^ and other by directly binding to protein or protein complexes. An example of the latter is the antisense lncRNA *ZFAS1*, which regulates Ca2+ flux in the cells by directly inhibiting the activity of the protein SERCA2a (sarcoplasmic reticulum Ca2+-ATPase 2a)^50^. Overexpression of *ZFAS1* altered Ca2+ flux leading to intracellular Ca2+ overload in CM. Other lncRNAs, such as the *Sirt1* antisense lncRNA binds to *Sirt1* mRNA 3′-UTR and subsequently increased *Sirt1* abundance^51^. Exemplified by these examples, *VHRT* repression that negatively regulates the CM contraction and structural gene program, prompts the need for further studies to understand how cytoplasmic *VHRT* supports and stabilizes the activity and abundance of cardiac contractility genes expression.

To date many genetic variants have been associated with cardiovascular disease^28^, but few have been linked through lncRNAs function. An example is the cardioprotective lncRNA *Myocardial infarction associated transcript* (*MIAT*). Six SNPs at the *MIAT* locus are associated with myocardial infarction (MI)^52^. More recently, three SNPs (e.g. rs7140721, rs3729829 and rs3729825) at the *Myosin Heavy Chain Associated RNA Transcript* (*MHRT)* locus are associated with increased risk of HF and prognosis^53^. Thus far, we have concluded, thorough integrative analysis of GWAS, cis-eQTL and RNA-seq from ∼ 31,5K sample patients^30^, that a series of SNPs in the *VHRT* locus also associated with heart disease. The SNPs rs6489844-G and rs11065780-C are both associated with decreased *VHRT* expression in human left ventricle and increased risk for HF. Moreover, *VHRT* is downregulated in failing hearts from patients with DCM, reflecting the link between *VHRT* and HF. Indeed downregulated contractile and calcium regulatory genes in failing DCM hearts, consistent with downregulated genes in VHRT KD, again concur with the possible mechanism mediated the contribution of VHRT to HF disease progression. The lack of species conservation for *VHRT* complicates *in-vivo* validation since a homologue for *VHRT* in rodents cannot be identified. Nevertheless, novel strategies to assess *VHRT* as a target for drug development are now attractive to pursue.

In conclusion, we have shown that *VHRT* is a crucial regulator for CM specification and lies upstream of key CM structural and contractile gene programs. Perturbation of *VHRT* results in the loss of CM contractile function. Moreover, concordant HF genome-wide analyses and human heart transcriptomic cross-comparisons motivate the need for follow up functional studies using models of heart disease.

## ACKNOWLEDGEMENTS

Singapore International Graduate Award (A.D.); EMBO long-term fellowship ‘EMBO ALTF 304-2012’ (M.K.B.J.). Funding from the Singapore National Medical Research Council (R.S.Y.F.) and A*STaR Biomedical Research Council (R.S.Y.F., L.B.). Our gratitude to Dr. William Sun and Dr. Kristy Purnamawati for the use of their multielectrode system.

## AUTHOR CONTRIBUTIONS

AD, experimental planning and execution, figure preparation, manuscript drafting and writing

WT, bioinformatics analysis, figure preparation

CGA-N, experimental planning, figure preparation, manuscript writing

BP, cell culture and differentiation optimization

AM, CRISPR/Cas9 assay and experiments

CB, generation of MYH6 reporter line

MA, cell culture and RNA preparation

VV, contractility-based assay

RH, preparation of buffers and experimental assistance

ZT, library preparation for sequencing

HW, experimental planning and analysis, figure preparation

BL, senior supervisor, funding support

AW, bioinformatics analysis, figure preparation

MJ, study design, experimental planning and execution, analysis, figure preparation, manuscript drafting and writing

RF, senior supervisor, funding support, manuscript writing

## METHODS AND MATERIALS

See Supplementary information online for a detailed description of all methods and materials.

## DETAILED MATERIAL AND METHODS

### Stem cell maintenance and differentiation

Human embryonic stem cell line H1 was maintained using mTeSR medium (Stemcell Technologies, 85850) on 1:200 growth factor-reduced Geltrex (Thermo fisher, A1413202) coated tissue culture plates and passaged regularly as cell aggregates every 4-5 days using ReLeSR (Stemcell Technologies, 05872), an enzyme-free dissociation reagent specific for human pluripotent stem cells). Two days prior to starting differentiation, cells were dissociated using Accutase (Stemcell Technologies, 07922) and seeded as single cells in Geltrex-coated 12-well plates (passage ratio 1:2, between 500’000-600’000 cells). Differentiation was performed following the published protocol by Lian et al.^21^ with modifications as follows. 10 µM of CHIR99021 (Stemcell Technologies, 72054) was added on day 0 and left for 24 hours followed by medium change. On day 3, 5uM IWP2 (Sigma Aldrich, I0536) was added using 50/50 mix of new fresh medium and conditioned medium collected from each well and left for 48 hours. Culture medium from day 0 until day 7 was RPMI1640 (HyClone, SH30027.01) plus B-27 serum-free supplement without insulin (Gibco, A1895601). From day 7 and onwards RPMI1640 with B-27 serum free supplement with insulin (Gibco, 17504044) was used and changed every 2-3 days.

### Generation of *MYH6*-GFP reporter line

EGFP cassette with kanamycin selection was inserted into BACs for *MYH6* (RP11-834J17, BacPac) immediately before the initiating Methionine (ATG) using recombineering (Quick & Easy BAC Modification Kit, KD-001, Gene Bridges GmbH). The Tol2 transposon cassette with Ampicilin selection mark was inserted into the loxp site of the BAC in the backbone using recombineering. Ten million H1 cells were cultured in CF1 conditioned medium (20% KO serum replacement, 1 mMl-glutamine, 1% non-essential amino acids, 0.1 mM 2-mercaptoethanol and 8 ng ml^−1^ of basic fibroblast growth factor in DMEM:F12) for 6 days and dissociated into single cells with TrypLE™ Express (Thermo Fisher, 12604021) and electroporated with 20 micrograms of Tol2 transposes and 100 micrograms of Tol2/EGFP modified Transposon-BACs. After electroporation, cells were re-suspended in conditioned medium with 10 μM ROCK inhibitor Y276329 (Y27632 (Stemcell technology, 72302). ROCK inhibitor was added for the first 48 hours after electroporation. Fifty μg/ml geneticin (Gibco, 10131035) was added for selection of positive clones 72 hours post-electroporation. Fourteen days later after drug selection, single colonies were picked into 24 well plates for expansion. Fluorescent *in situ* hybridyzation (FISH) using non-modified BACs as probes was carried out to validate the incorporation of BAC construct into genome of ES cells (Cytogenetics Services, Genome Institute of Singapore). Karyotyping was performed to confirm a normal chromosome pattern.

### Tri-lineage differentiation

STEMdiff™ Trilineage Differentiation Kit (Stemcell Technologies, 05230) was used to differentiate hESC-MYH6 reporter into the three germ layers: ectoderm (*NESTIN* and *PAX6*), mesoderm (*TBXT* and *CXCR4*) and endoderm (*SOX17*, *FOXA2* and *CXCR4*) following manufacturer’s instructions. Differentiation was performed on a 24-well plate. At the end of the differentiation RNA was extracted and RTqPCR was run for respective lineage markers.

### Immunofluorescent staining

Cells were fixed in 3.7% formaldehyde for 15 min at room temperature and stored in DPBS. They were permeabilized in 0.2% Triton X-100 for 15 min followed by a pre-blocking step with 2% BSA for 20 min. Primary antibody incubation was performed in DPBS + 10% goat serum (except for Nkx2.5 for which donkey serum was used) overnight at 4 degree and secondary antibody incubation for 2 hours at room temperature. DAPI was included during the final washing step. Antibodies used were cardiac troponin T (Lab Vision, ms-295-P0, mouse, 1:500 dilution), α-actinin (Sigma Aldrich, A7811, mouse, 1:1000 dilution), Nkx2.5 (Santa Cruz, sc-8697, goat, 1:200 dilution), MLC2v (ProteinTech, 10906-1-AP, rabbit, 1:200 dilution), ryanodine receptor (Thermo Fisher, MA3-925, mouse, 1:200), alexa fluor 594 goat-anti-mouse (Thermo Fisher, A-11020), alexa fluor 546 goat-anti-rabbit (Thermo Fisher, A-1107), alexa fluor 647 mouse-anti-goat (Thermo Fisher, A-21237).To improve signal and avoid unspecific binding, MLC2v was stained using cells that were fixed in ice cold methanol for 5 mins at 4 °C.

### RNA and DNA isolation

RNA was extracted using Direct-zol™ RNA MiniPrep Kit (Zymo, R2060). Cells were directly lysed using Trizol reagent (Thermo Fisher, 15596026). DNA was purified using PureLink Genomic DNA Mini Kit (Thermo Fisher, K182001). All experiments were performed following the manufacturer’s instructions.

### PCR and reverse transcription quantitative PCR (RT-qPCR)

DNA or plasmid vector were PCR amplified using Q5 High-Fidelity 2X Master Mix (Bio Labs, M0492S) and target specific primers (IDT) following manufacturer’s instructions. RNA (50-500 ng) was reverse transcribed to cDNA using qScript Flex cDNA Kit (Quantabio, 95049-025) with a combination of random primers and oligo (dT). The remaining cDNA (1:10) was mixed with PerfeCTa SYBR Green FastMix, low ROX (Quantabio, 95074-05K) and specific primers on a 384-well plate. Real time qPCR was run using ViiA 7 Real-Time PCR System (Thermo Fisher). Average Cq was recorded and ΔΔCq method was used to calculate relative gene expression changes. Expression levels of genes were normalized against two housekeeping genes, *GAPDH* and *PPIA*. List of primers and sequences used in this study are listed on **Supplementary Table 11.**

### Single cells RNA-seq

#### Single cell preparation and cDNA preparation for RNA-seq

hESC-CM were dissociated into less dense monolayers > 3 days prior to RNA isolation by pre-treatment with 1 mg/ml collagenase IV for 1 hour at 37 °C followed by 5 minutes treatment with Accutase (Stemcell Technologies, 07922). Immediately prior to single cell capture, cells were dissociated using Accutase (Stemcell Technologies, 07922) for 4-5 minutes at 37 °C and re-suspended with a P1000 pipette to facilitate proper single cell formation. Cells were filtered using a 40 µM filter and kept in suspension in RPMI1640 with B-27 supplement and 5 uM ROCK inhibitor Y276329 (Stemcell technologies, 72302) inhibitor on ice and counted using the TC20 automated cell counter from Biorad. The cell size ranged between 16-22 µM and cell viability was above 95%. Fluidigm microfluidic C1 system was used for automated cell capture, conversion of polyA+ RNA into cDNA and a universal amplification step. After priming of the chip following Fluidigm manual, cells were loaded onto large sized C1 IFC chips (Fluidigm, 100-5761) at a concentration of 400 cells/ul (total volume of 20 ul with a 3:2 ratio of cell suspension:C1 suspension reagent). The mRNA Seq: Cell Load script was run followed by manual inspection to assess cell capture success. Pictures were taken from all 96 capture sites and a note was made whether 0, 1 or more cells were captured per site and if the captured cell was GFP negative or positive. Cell lysis, reverse transcription and PCR amplification was run overnight (using reagents from the SMARTer Ultra Low RNA Kit from ClonTech, 634833) and the cDNA products harvested and diluted in 10 µl of C1 DNA Dilution Reagent (from the C1 Reagent Kit for mRNA seq from Fluidigm, 100-6201) the next morning.

#### Library preparation and Quality Control

cDNA concentration was measured using the Quant-iT™ PicoGreen® dsDNA Assay Kit (LifeTech, P11496) and ranged from 0.1-1 ng/ul per cell. 3µl of cDNA was used for library preparation with the Nextera XT DNA Sample Preparation Kit (Illumina, FC-131-1096). Each library was barcoded and multiplexed 15 libraries into one sequencing lane. A total of 71 GFP positive (CM) and 31 GFP negative (non-CM) cells were sequenced on an Illumina HiSeq 2000 sequencer in 4 different sequencing runs. From the 71 GFP positive cells, 4 were duplicated in one run to control for technical variability. 3 samples (one from each time point) that were first ran in individual runs were subsequently re-sequenced together in the fourth run to control for batch effects. Three samples (one from each time point) also underwent a new round of library preparation and were sequenced together with the original library prepared sample to assess the influence of the library preparation step on the outcome.

#### Raw reads filtering, alignment, quality control and gene expression estimation

Illumina CASAVA version 1.8.2 was used to perform demultiplexing and generate FASTQ file. Quality of sequencing reads was examined with FASTQC^1^. Paired-end reads were aligned to human genome (*Homo Sapiens*) hg19 assembly using state-of-the-art mapping tool, Tophat2^2^ (version 2.0.11.Linux_x86_64) with Bowtie2 (version 2.2.1.0). The parameters that were set for Tophat2 alignment were –min-anchor 8, -- splice-mismatches 1, --min-intron-length 50, --max-intron-length 500000, --min-isoform-fraction 0.15, -- max-multihits 1, --segment-length 25, --segment-mismatches 2, --min-coverage-intron 50, --max-coverage-intron 20000,--min-segment-intron 50, --max-segment-intron 50000, --read-mismatches 3, --read-gap-length 3, --read-edit-dist 3, --read-realign-edit-dist 0, --max-insertion-length 3, --max-deletion-length 3, -- mate-inner-dist 200, --mate-std-dev 20, --no-coverage-search and –library-type fr-unstranded. We also suppliedTophat2 with hg19 GENCODE (version 19) transcriptome annotation to Tophat2 with –G option. RSeQC (RNA-seq Quality Control Package version 2.6.1) was used to inspect sequence quality, transcript integrity number (TIN) which is analogous to RIN (RNA integrity number), duplication rate and mapped reads distribution^3^. Reads mapped to mitochondrial genes and ribosomal genes were also calculated using htseq-count (version 0.6.0).

Cuffdiff (version 2.2.1, with boost version 104700) was used to compute relative gene expression level in human genome (hg19) GENCODE (version 19) transcriptome annotating tf file with parameters: --library-norm-method classic-fpkm, --compatible-hits-norm, --frag-bias-correct and --max-frag-multihits 1. Ribosomal RNA and tRNA were masked in the gene expression level calculation. Gene expression level was reported in fragments per kilobase per million mapped reads (FPKM)^4^. We assumed genes with FPKM lower than 1 to be non-expressing. Only genes expressed with FPKM ≥ 1 in at least 2 samples were considered for downstream analyses.

#### Saturation Analysis

Using samtools version 0.1.19, saturation analysis was performed by randomly subsampling different number of reads from each sample, and re-calculating gene FPKM. The process of subsampling was repeated until there were at least 5 subsampled datasets per point, at library size of (0.1M, 0.5M, 1M, 2M, 3M, 4M, 5M). For each point, the average number of genes with FPKM greater than or equal to 1 was plotted.

#### Network construction and module detection using R package Weighted Correlation Gene Network Analysis (WCGNA)

Using WCGNA, we constructed a signed weighted correlation network by computing pairwise correlations, s between all genes across all single-cell RNA-seq samples^5^. Next, we chose soft thresholding power (β=3), in constructing an adjacency matrix using the formula, a_ij_ = (0.5 + 0.5 × s_ij_)^β^, where a_ij_ is defined as weighted correlation and s_ij_ is defined coefficient correlation between betweengene_i_ and gene_j_. We choose the power (β=3), which was the lowest power for which the scale-free topology fit index curve flattens out upon reaching a high value of 0.98. Using the adjacency matrix computed in the previous step, topological overlap was calculated to measure the network interconnectedness in a robust and biological meaningful way. The topological overlap was utilized to group highly correlated genes together using average linkage hierarchical clustering.

Modules were defined as the branches obtained by cutting the hierarchal tree using Dynamic Hybrid Tree Cut algorithm^6^. We defined the first principle component of a module as module eigengene, which was representative of the expression profile in each module. Genes in each module were removed if the correlation between the gene and module eigengene (kME) was less than 0.3. If a detected module did not have at least 30 genes with eigengene connectivity (kME) at least 0.5, the module was disbanded and its genes were unlabeled and returned to the pool of genes waiting for module detection. Modules whose eigengenes were highly correlated (correlation above 0.75) were merged. Construction of signed gene network and identification of modules was performed using R function, blockwise Modules with following parameters: soft thresholding power = 3, minimum module size = 30, mergeCutHeight = 0.25, corType = “Pearson”, networkType = “signed”, TOMType = “signed”, minCoreKME = 0.5, minKMEtoStay = 0.3. To identify modules that were significantly correlating with subgroups, we computed correlation of eigengenes of each module with cells in W02, W06 and W12. We picked modules which showed high correlation with each week and sub-population of cells for W12. In addition, we also computed Gene Significance (GS), which was defined as the correlation of each gene with each group of interest^5^. We also calculated module membership, which was defined as the correlation between module eigengene and gene expression profile. GS and MM were important because it helped in the identification of genes with high significance for each week of development and high module membership in each week-/subtype-specific modules. Module membership was reported to be highly correlated to the intramodular connectivity, k_ME_. Highly connected intramodular hub genes was observed to have high module membership values to the respective module.

### Bulk RNA seq analysis

Total RNA sequencing was performed using Truseq Stranded Total RNA Library Prep kit (Illumina, RS-122-2201), which uses Ribo-Zero to remove cytoplasmic rRNA from total RNA. Remaining intact RNA is fragmented using a chemical mix, followed by first- and second-strand cDNA synthesis using random hexamer primers. “End-repaired” fragments were ligated with unique Illumina adapters. All individually indexed samples were subsequently pooled together and multiplexed for sequencing. Libraries were sequenced using the Illumina Hiseq 2000 sequencing system and paired-end 101 bp reads were generated for analysis. Quality control, gene alignment and gene expression estimation were performed as described in the single cell sequencing method.

### GWAS and eQTL analysis and statistics

Genome-wide association studies (GWAS) summary statistics were accessed from the Cardiovascular Disease Knowledge Portal (CVDKP)^7^, and included the following data sets: All-cause Heart Failure and Nonischemic Cardiomyopathy HRC GWAS in UK Biobank (394156 samples, European ancestry)^8^, CARDIoGRAMplusC4D GWAS (184305 samples, mixed ancestry)^9^, CAD exome chip analysis (120575 samples, mixed ancestry)^10^ and 2018 AF HRC GWAS (588190 samples, mixed ancestry)^11^. Significant VHRT cis-eQTLs (FDR<0.05, +/-1Mb window around the VHRT TSS) precalculated from genotype and RNA sequencing data from human heart left ventricular samples (n=386) were extracted from the Genotype-Tissue Expression (GTEx) Portal^12^. VHRT locus plots depicting GWAS and eQTL data were created using LocusZoom^13^ on genome build GRCh37/hg19 and LD population 1000 Genomes Phase 3, EUR.

### Cross-interrogation of *VHRT* KD transcriptome with human DCM date sets

Significant DE genes in GapmeR#1 VHRT KD and GapmeR#2 VHRT KD vs CTRL (N=1,100; 573 down- and 527 upregulated at adjP-value<0.05, FC>|0.5| in both) were compared with public DCM transcriptomics study (GEO accession: GSE141910). This study comprises RNA sequencing datasets of 61 left ventricle explants (Idiopathic Dilated Cardiomyopathy and Non-Failing) from the MAGNet study (Myocardial Applied Genomics Network). Differential expression analysis was conducted on the DCM versus control samples using R package EdgeR, which applies generalized linear models (glm) on normalized RNA-seq gene count. Enrichment for pathway and molecular functions was then performed on the DE subgroup based on functional analysis using Enricher tools^14^.

### *VHRT* CRISPR/Cas9 knockout

Plasmid pMIA3 (Addgene plasmid # 109399; http://n2t.net/addgene:109399; RRID: Addgene_109399) was used for CRISPR/Cas9 mediated KO. pMIA3 was generated and previously described by Dr. Matias Autio^15^. Two KO hESC lines were generated using dual single guide RNA (sgRNAs) to remove part of VHRT. Single guide RNA sequence were designed using cloud based software tools for digital DNA sequence editing Benchling^16^ and CRISPOR^17, 18^. Schematic representation of the pairs of sgRNAs and genomic regions to be removed are shown in the figure 5A. 10 µM sense and anti-sense oligonucleotides were ordered **(Supplemental Table 11)** and annealed to generate the 20 nucleotide spacer that defines the genomic target to be modified (VHRT 5’end and 3’end). pMIA3 was digested with BsmBI and sgRNA spacers cloned after human U6 promoter with T4 DNA ligase (New England Biolabs) following manufacturer’s instruction. Ligated constructs underwent transformation using RapidTrans™ Chemically Competent Cells (Active motif). Plasmid was extracted from bacterial culture, purified and Sanger sequenced to confirm successful cloning. Prior to hESC targeting, cutting efficiency of pMIA3-sgRNAs plasmids were tested on HEK293T using the EGxxFP plasmid (pCAG-EGxxFP was a gift from Masahito Ikawa, Addgene plasmid # 50716). Best combination of sgRNAs was then used for final targeting of hESCs. The best pair of guides were cloned into a single pMIA3 plasmid digested with NheI & XbaI, using NEBuilder isothermal assembly (New England Biolabs), according to manufacturer’s instructions to make the final pMIA3 dual sgRNA plasmid. Primers for isothermal assembly are listed on **Supplemental Table 11.** Human ESCs were dissociated with Accutase (Stemcell Technologies, 07922) and ∼1.5×10^6^ cells were re-suspended in 100 μl P3 Primary Cell kit solution from Lonza (V4XP-3024) and mixed with 10 μg pMIA3dual sgRNA plasmid. To transfect hESC, nucleofection was performed using program CM-113 on the 4D-Nucleofector System (Lonza). Cells were then plated into Geltrex coated 6-well plate with mTeSR medium (Stemcell Technologies, 85850) containing 5 μM Y-27632. After 2 days in culture, cells were dissociated, FACS sorted for RFP positive cells and collected into a tube containing TeSR medium (Stemcell Technologies, 85850) with 5 μM Y-27632. 500 to 2000 cells were plated into wells of 6-well plate containing the above media. Single cell clones were monitored and upon sizeable growth, colonies were picked and passaged. Genomic DNA was extracted for genotyping and screened for successful KO. Location of primer pairs used for the genotyping are represented in **Supplemental Figure 7A-C** and primer sequences are listed in **Supplemental Table 11**. RT-qPCR was performed to validated KD of *VHRT* transcript.

### Cellular fractionation

An average of 10^7^ hESC-CMs were dissociated into single cell suspension as described above. RNA was extracted from nuclear and cytoplasmic fractions using a rapid, phenol-free, small scale protein and RNA isolation system PARIS kit (Thermo Fisher, AM1921) following manufacturer’s instructions. During the fractionation procedure, Trypan blue solution was used to monitor proper separation of the cytoplasmic and nuclear fractions **(Supplemental Figure 5B)**. To remove traces of genomic DNA, eluted RNA was treated with DNA-free™ DNase Treatment and Removal Reagents (Thermo Fisher, AM1906) following manufacturer’s instructions. RNA concentration and purity were measured using NanoDrop. An Agilent RNA 6000 Pico kit (5067-1513) was run to observe the RIN value, distribution and intensity of the signal of all RNAs, up to about 7000 nt, hence including the distinctive peak for tRNA, 18S and 28S. *VHRT* compartmentalization was detected with RT-qPCR along with cytoplasmic markers (*GAPDH* and 18S ribosomal RNA) and nuclear markers (*U6* and *MALAT1*).

### *In-vitro* translation

Protein coding potential was assessed via web-based prediction algorithm (CPC: coding potential calculator). An open reading frame (ORF) was identified on *VHRT* isoform 6. 1x FLAG Tag (motif: DYKDDDDK) was inserted either at N-terminus (after the START codon) or the C-terminus (before STOP codon) of the ORF. Plasmids were transfected in HEK293T cells. Successful transfection was confirmed using a plasmid control containing sequence for blue fluorescent protein (BFP: ∼29 kDa) tagged with FLAG at the C-terminus. 48 h after transfection, cells were lysed for protein extraction and Western blot. Protein analysis was performed using pre-casted gradient gel 4-20% mini-PROTEAN TGX Stain-Free (Bio-rad, 4568096). 10 to100 μg of protein was loaded onto the gels alongside a protein ladder (1st Base, Asia). The electrophoresis setting for initial running for 20 mins at 50 V and subsequently 100 V for 60-90 mins. The running buffer contained 25 mM Tris base, 192 mM glycine and 0.1 % SDS. Protein transfer was performed on 0.22 μm polyvinylidene diflouride (PVDF, BioRad, 1620260) membrane at 100 V for 1 h in transfer buffer (25 mM Tris base, 192 mM glycine, 20 % methanol). After transfer, the membrane was blocked in 5 % BSA/TBS-T (20 mM Tris-HCl pH 7.6, 140 mM NaCl supplemented with 0.1 % Tween 20, Merck, P1379) for 1 h on a rocking platform at room temperature (RT). Membranes were subsequently incubated with primary anti-mouse FLAG antibody (Sigma Aldrich, B3111) diluted in 5 % BSA/T-BST blocking buffer overnight at 4°C. Membranes were washed 3 times in TBS-T for 5 min. Horse radish peroxidase (HRP)-conjugated secondary antibody (Thermo Fisher, MA5-15367) was diluted in 5 % BSA/T-BST at 1:10 000 dilution and added to the membrane for 1h at RT. After membranes washing with TBS-T, chemiluminescence solution (West Femto, Thermo Fisher, 32106) was added to the membrane following the manufacturer’s instructions and exposed to chemidoc (Biorad) for protein exposure.

### Human heart tissue collection

Gene expression level was assessed by qPCR from explanted hearts of patients undergoing transplantation for end-stage heart failure. All donors provided written informed consent and samples were fully anonymized. 30-60 mg of cardiac tissue (Ventricle Tissue, VT and Atrial Tissue, AT) were mechanically powdered on liquid nitrogen and further disrupted in 350 µl RLT buffer at room temperature using a Tissue Lyzer device (Qiagen, Hilden, Germany; 2 min, 30 Hz). DNA extraction was performed with the RNeasy Plus kit (Qiagen) according to the manufacturer’s instructions. 100-250 ng of RNA were reversely transcribed with the High Capacity cDNA Reverse Transcription kit (Applied Biosystems). qPCR was carried out on an ABI Prism 7900 HT Sequence Detection System (Applied Biosystems). Transcript abundance was calculated using the the delta-delta-CT method with GAPDH as reference transcript.

### VHRT knockdown

#### GapmeR transfection

Anti-sense LNA - GapmeR –based technology (Qiagen) was used for gene KD. With the use of a web-based algorithm provided by Qiagen, GapmeRs were designed to target *VHRT* transcript (RP1-46F2.2, hg19 or LINC01405, hg38). 100 nM of each GapmeR was transfected using 6 ul Lipofectamine 2000 reagent (Invitrogen, 11668030) in a total volume of 1mL hESC-CM media (RPMI 1640 with B27 supplement with insulin, Thermo Fisher Scientific, A1895601) in a 12-well plate with 85-90% cell confluency (n= 4 biological replicates). Negative A–GapmeR was transfected as negative control. Sequence of GapmeRs are listed in **Supplemental Table 11.**

Fresh and new media was replaced after 48 h. Day 4 after transfection, RNA was purified and relative expression of *VHRT* and other important cardiac genes (*MYH6, MYH7, RYR2, ATP2A2, SCN5A* and *PLN*) were assayed via RT-qPCR in each group. Expression levels were normalized to housekeeping genes *GAPDH* and *PPIA*. % KD was calculated relative to Negative A-GapmeR and genes that had significant changes in expression (*p*-value ≤ 0.05, Student’s t test) were considered statistically significant.

#### SiRNA transfection

*In-vitro* iCell cardiomyocytes (CDI, 01434) were maintained in culture according manufacturer instruction. SiRNA for *VHRT* was purchased from Riboxxx life science (lot 2017d0113) and for *ATP2A2* from Dharmacon (ON TARGETplus SMARTpool siRNA). They were dissolved in H_2_O to a stock concentration of 10 µM. 10 nM siRNA was transfected using 0.1 uL Lipofectamine RNAiMax per well. Media was changed after 24h and cells were used in the corresponding assay after 72h. RNA was extracted using the RNeasy micro kit (Qiagen, 74004) in the Qiacube instrument. cDNA was prepared using reverse transcription according to High-Capacity RNA-to-cDNA™ Kit (Thermo Fisher, 4387406). qPCR was performed using TaqMan Fast Advanced Master mix (Thermo Fisher) together with specific gene expression assays Hs04405003_m1 (*RP1-46F2.2*) and housekeeping gene (*GAPDH*). Quantitative RT-PCR was performed with QuantStudio™ 7 Flex Real-Time PCR instrument (Thermo Fisher) and data was recorded during 40 PCR cycles and normalized against untreated samples. Sequence of siRNAs are listed in **Supplemental Table 11.**

### Fluorescence-activated cell sorting (FACS) of hESC-CM

Cells were sorted using BD FACSAria II 5 laser from Flow Cytometry Core or (A*STAR, Singapore Immunology Network). Prior dissociation, cells were cultured with corresponding media containing 5uM ROCK inhibitor (Y27632, Stemcell Technologies, 72302) for 30 min at 37○C. Media was aspirated and cells were treated with Accutase (Stemcell Technologies, 07922) (1mg/mL) for 5-8 min at 37○C. For hESCs, Accutase (Stemcell Technologies, 07922) was neutralized using mTeSR medium (Stemcell Technologies, 85850) whereas passaging media was used for hESC-CMs. Cells were collected in 15 ml tube and spin at 300 g for 5 min. Human ESC were re-suspended in DPBS and hESC-CMs in FACS buffer (DPBS w/o Ca and w/o Mg, 2% HI-FB, 11550356, + 1mM EDTA, 5uM Y27632) and pipetted through 40 um cell strainer to avoid cell clumps. Due to their increase in cell size over time, older hESC-CMs were pipetted through a 70um cell strainer instead. During FACS, cells were collected in 2mL tube containing corresponding media and re-plated on a new coated plate (1:200 Geltrex, if hESC or 0.1% gelatin, if hESC-CM). To avoid contamination 100x Penicillin-Streptomycin mixture was added at each step and sample were kept on ice at all time to reduce cell death.

### Electrophysiological assays

#### Multielectrode array recordings

MEA chips were coated with 0.1% gelatin and hESC-CM (8 weeks after differentiation) were seeded on the chip at least 4 days prior to the recording to allow for proper attachment and coupling. Field potentials were recorded from spontaneously beating cells using a USB-MEA60-Up-BC-System (Multichannel systems) and analyzed using MC_Rack. Recordings were made in regular RPMI-1640 (Thermo Fisher, 11875093) +B-27 supplement media (Thermo Fisher, A3582801) with our without the addition of 100 µM noradrenaline.

#### Contractility and electrophysiology recordings

The Cardioexcyte 96 instrument (Nanion) was used to record and analyze impedance-based contractility. The stim plates were coated with fibronectin for >1h in the incubator. iCell cardiomyocytes (01434 from CDI) were thawed and plated at 20k cells/well following recommendation from the manufacturer. Experiments were performed from day 12 post-plating and onwards. At least 5 replicates were used for each condition. *ATP2A2* was used as positive control and non-targeted siRNA as well as untreated cells as negative controls. Readings for base impedance were measured before and after treatment with siRNAs, both for spontaneously beating and electrically paced cells. The data was analyzed using the data control software from Nanion. One-way ANOVA, multiple comparisons, compared to control siRNA.

#### Action potential (AP) recordings

Whole cell configuration of the patch-clamp technique was used to measure AP on two different groups of cells, Negative A-GapmeR and *VHRT* KD GapmeR. Both groups were co-transfected with BLOCK-iT™ Alexa Fluor ^®^ Red Fluorescent Control (Invitrogen), therefore only cells marked in red and green (*MYH6*-GFP reporter) were used to record AP. The signal was amplified using an axon 700B patch clamp amplifier (axon instrument, Sunnyvale, USA) and low-pass filtered at 5 kHz. Patch pipettes were fabricated from glass capillaries (O.D, 1.5mm; I.D, 0.9mm) using a Sutter P-97 microelectrode puller (Novato, CA, USA) and the tips were heat polished with a microforge (NARISHIGE MF-900) to gain a resistance of 2-4 MΩ. The electrical signals were sampled at 2.5-10 kHz and filtered at 2 kHz using a low-pass filter. Data acquisition was achieved using the Digidata 1440A (axon instrument). Data analysis and fit were performed using clamp fit 10.2 (axon instrument) and Origin 7.0 software (Origin Lab Corporation). A pClamp software (Version8.1; Axcon Instrument) was used to generate voltage-pulse protocols, acquire and analyze data. APs were recorded under current-clamp mode at 35 °C. Pipette solution contained (in mM): KCl 130, NaCl 5, MgCl2 1, MgATP 3, EGTA 10, and HEPES 10, with pH adjusted to 7.2 with KOH. Extracellular solution (Tyrode’s solution) containing contained (in mM) NaCl 140, KCl 5.4, CaCl2 1.8, MgCl2 1, glucose 10, HEPES 10, with pH adjusted to 7.4 with NaOH. The parameters of APs include AP durations (APD) at 20%, 50% and 90% of repolarization (APD20, APD50, and APD90), AP amplitude (APA), maximal diastolic potential (MDP), and beating frequency (for spontaneous contractions) were analyzed.

### Statistics

Excluding the NGS seq results, all gene expression analysis was performed using Excel (Microsoft office). Unless stated otherwise, data are represented as mean ± standard error of the mean (s.e.m). Statistical significance between two groups was done using student’s t. P-values of ≤ 0.05 were considered as significant, unless otherwise indicated.

## SUPPLEMENTARY FIGURE LEGENDS

**Supplementary Figure 1.**
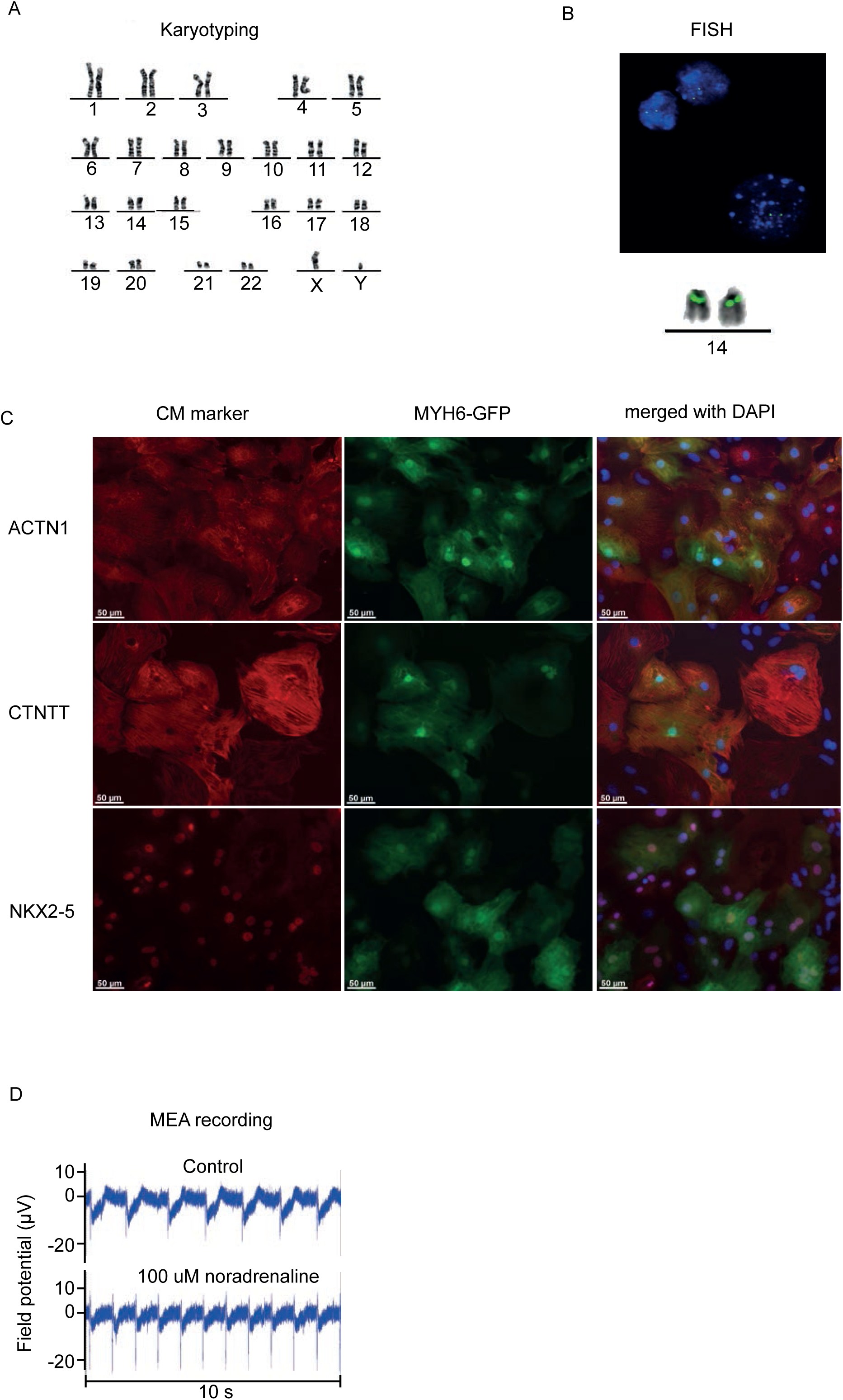
**(A),** Karyotyping of the *MYH6*-GFP reporter cell line shows a normal karyotype. **(B),** FISH identified 1 signal incorporated on each allele of chromosome 14. **(C),** Immunofluorescence confirms protein expression of α-actinin (ACTN1), cardiac troponin T (cTnT) and NKX2-5 in hESC-derived cardiomyocytes (CMs) indicating high differentiation efficiency. Striated patterns with ACTN1and cTnT are typical for cardiac muscle. Overlap with GFP shows that all GFP positive cells expressed all three cardiac proteins. **(D),** A multielectrode array was used to record field potentials from hESC-CM under baseline conditions and upon administration of adrenergic stimulant noradrenaline. Physiological field potentials are as displayed showing that noradrenaline induced an increase in beating frequency, increased field potential amplitude and a shortening of the field potential duration.

**Supplementary Figure 2.**
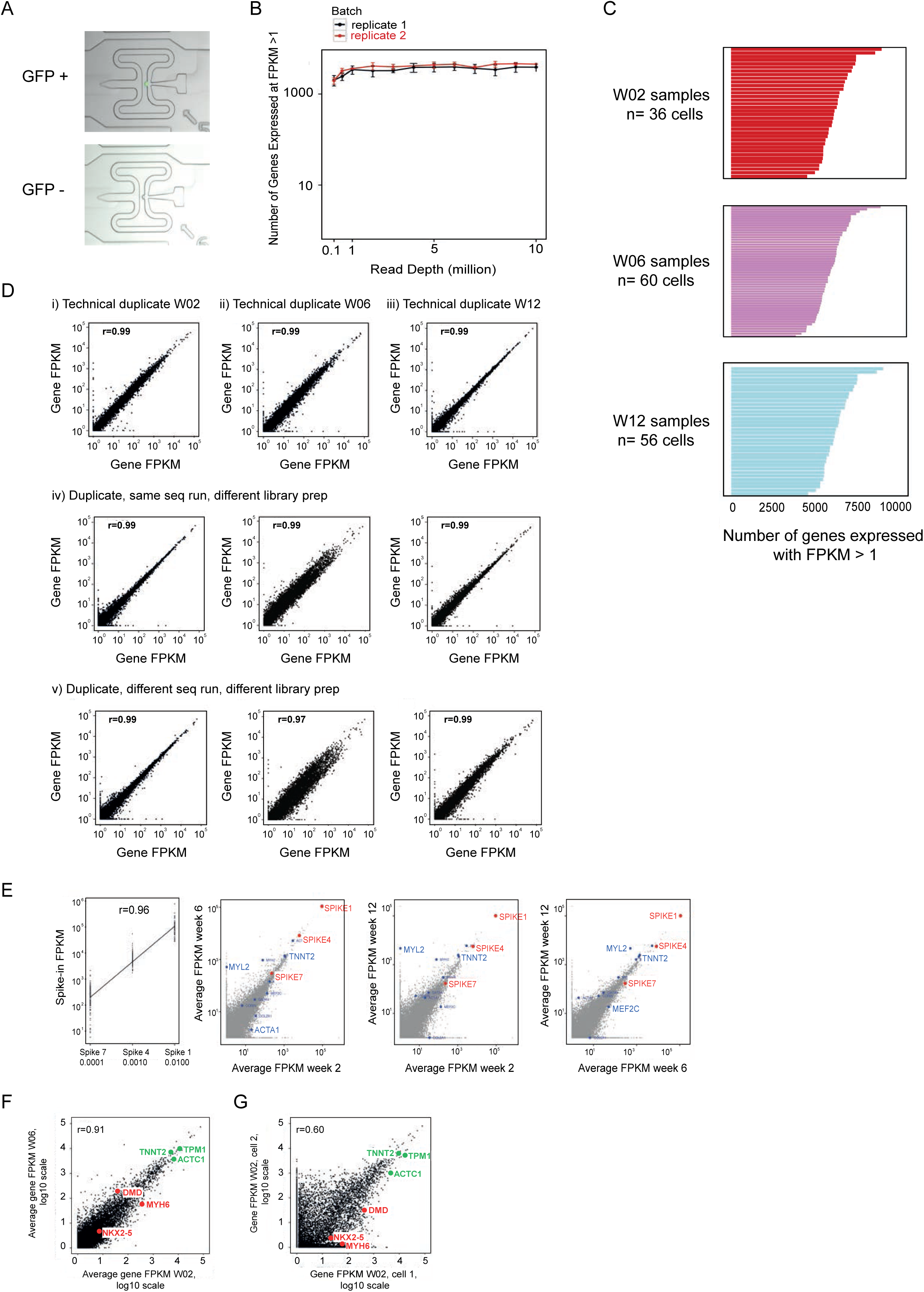
**(A),** Single cells captured in the Fluidigm C1 microfluidic system. Top, GFP positive (GFP+) and bottom, GFP negative (GFP-) cells, respectively. **(B),** Saturation plot for all genes expressed at FPKM ≥ 1 shows that for all samples, the maximum number of expressed genes was attained by around 1 million sequencing reads. **(C),** Histoplot showing the number of expressed genes (FPKM >1) detected in each single cell sample at the different time points. **(D),** Scatter plots for correlations between technical repeats show strong reproducibility and excludes any predominant effect of technical noise or batch effect. First, three libraries (one from each time point) were re-sequenced together in one sequencing lane and the results compared to the original sequencing of the same library (i-iii). Second, cDNA from previously sequenced samples were subjected to a new round of library preparation and re-sequenced together with initially prepared libraries (iv). Third, freshly prepared and sequenced libraries were compared to sequencing from the initial corresponding libraries (v). **(E),** Spike-in concentration vs FPKM values show strong linear correlation (r=0.96). Spike-in values in 2 weeks vs 6 weeks samples, 2 weeks vs 12 weeks, and 6 weeks vs 12 weeks show high technical consistency. **(F),** Scatterplot comparing averaged gene expression in W02 to W06 cells, showing a high correlation r value (0.91), the expected lack of differential expression for stable cardiac genes (green dots: *TNNT2*, *ACTC1*, *TPM1*), but indistinct differential expression of variable cardiac genes (red dots: *MYH6*, *NKX2*-5, *DMD*) at this averaged pooled level of analysis. **(G),** Scatterplot comparing gene expression in a single W06 cell to another single W06 cell, now showing a lower correlation r value (0.6) than (E), the same lack of differential expression of stable cardiac genes (green dots), but instead distinct differential expression of the 3 variable cardiac genes (red dots).

**Supplementary Figure 3.**
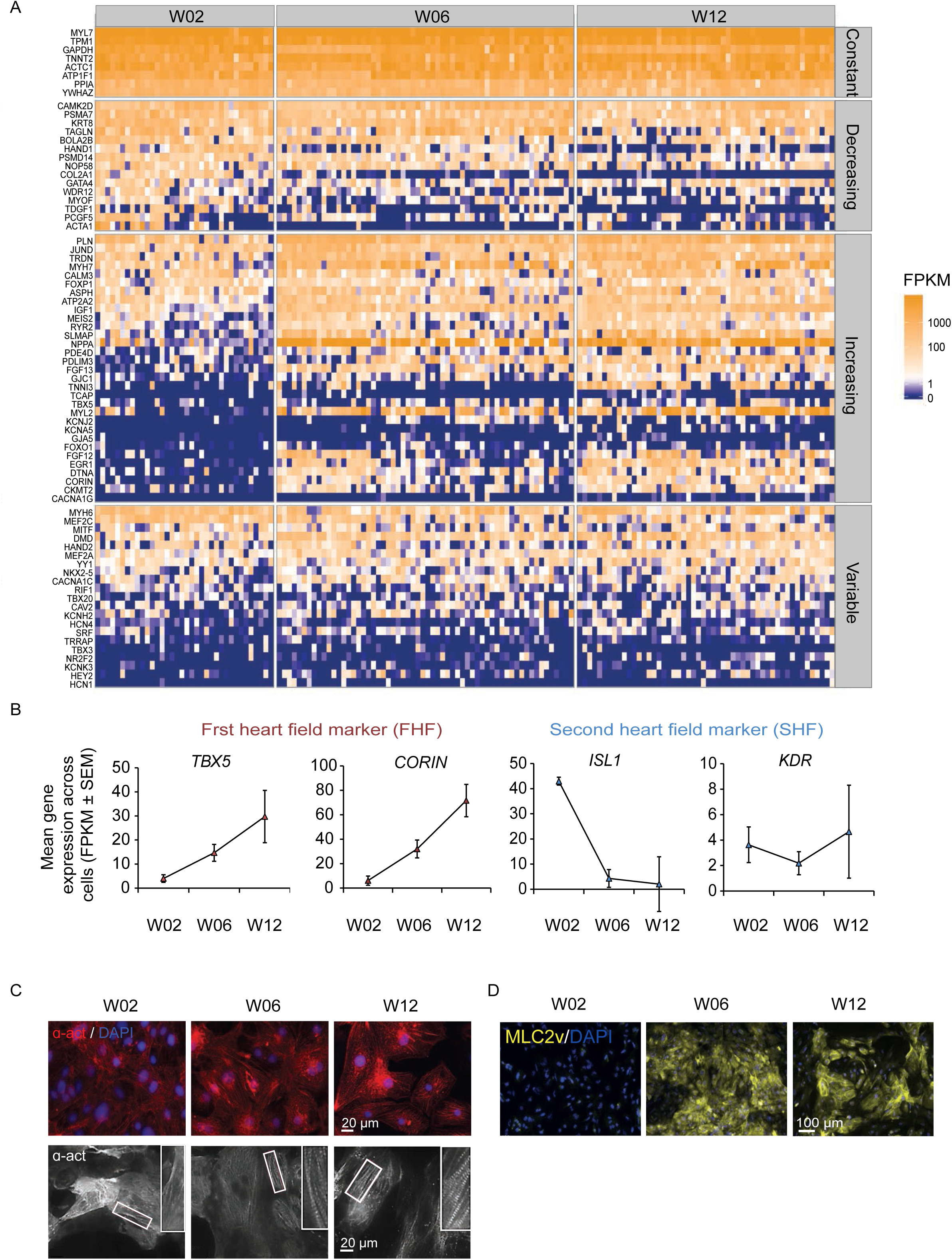
**(A),** Heatmap of cardiac genes that have either (a) constantly high expression in all cells at all-time points, (b) decreasing, or (c) increasing abundance in expression over time, or (d) highly variable expression across all time points. **(B),** Line plots showing mean gene expression across cells for first and second heart field markers. Data are represented as mean FPKM ± s.e.m. **(C),** Immunostaining confirms protein expression at all-time points for cardiac alpha actinin (α-act, top panel) and displaying sarcomere organization (α-act, bottom panel). **(D),** Immunostaining showing MLC2v protein expression appearing only from W06.

**Supplementary Figure 4.**
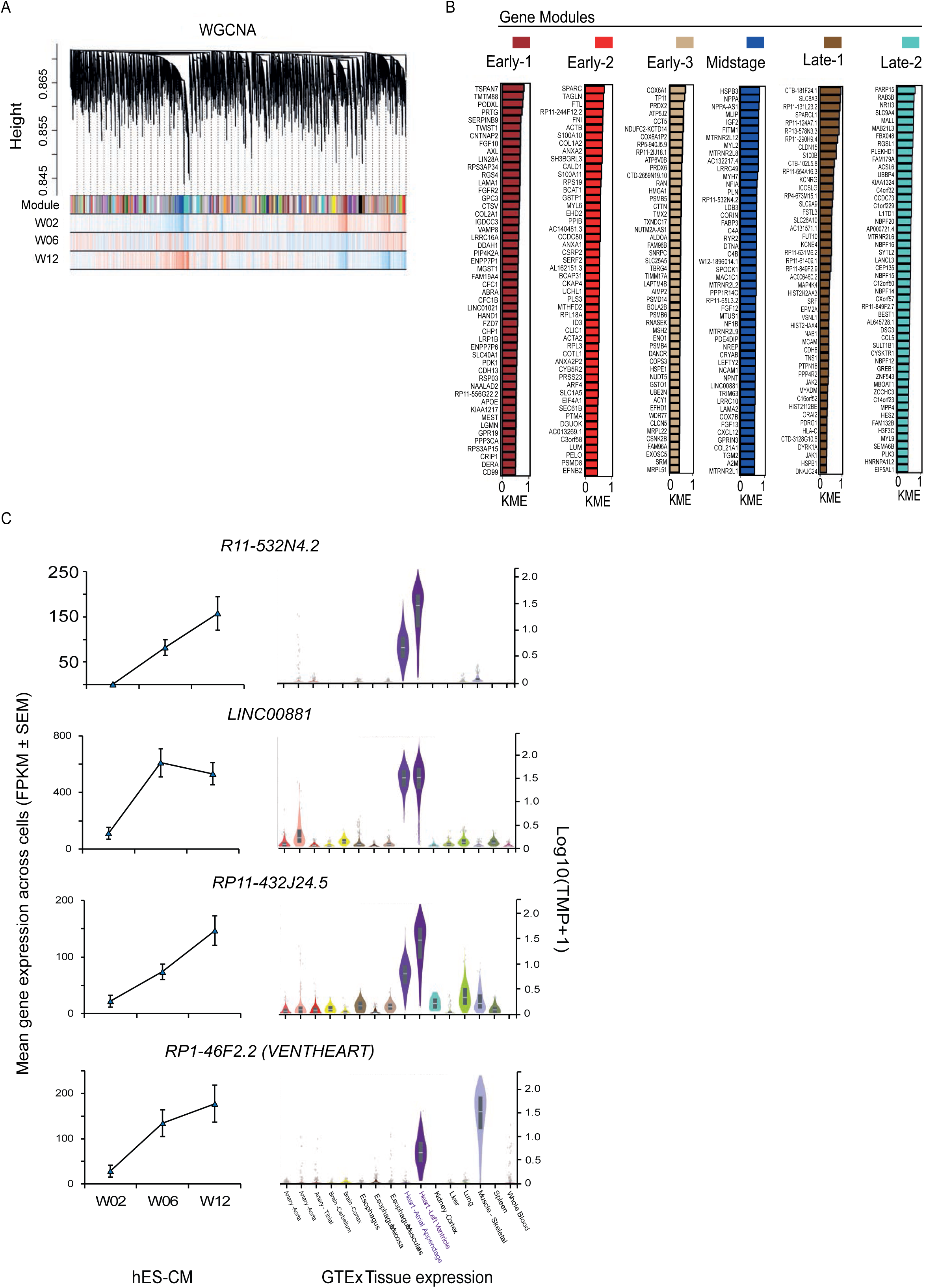
**(A),** Module dendrogram showing gene-network modules that were derived using weighted gene co-expression network analysis (WGCNA). Each vertical black line represents a uniquely expressed gene, grouped according to co-expression gene modules (colored bars). Correlation between gene modules and the time points are represented in the 3 horizontal bars (blue for negative-correlation and red for positive-correlation). Six gene modules differed significantly between at least 2 of the time points. **(B),** Top 50 expressed genes in each of the 6 modules. KME refers to module membership of each gene where values closest to 1 indicate genes that are most highly co-regulated with the other genes in the same module. **(C),** Expression of 4 annotated lncRNAs associated with blue (midstage) module are represented. Line plots showing increased mean lncRNA expression across single cells in W02, W06 and W12 hESC-CM. Values are represented as mean gene FPKM ± s.e.m. Right panel shows violin plot with expression profile across different tissues from GTEx data set. Data are represented as log Transcripts Per Kilobase Million (TMP).

**Supplementary Figure 5.**
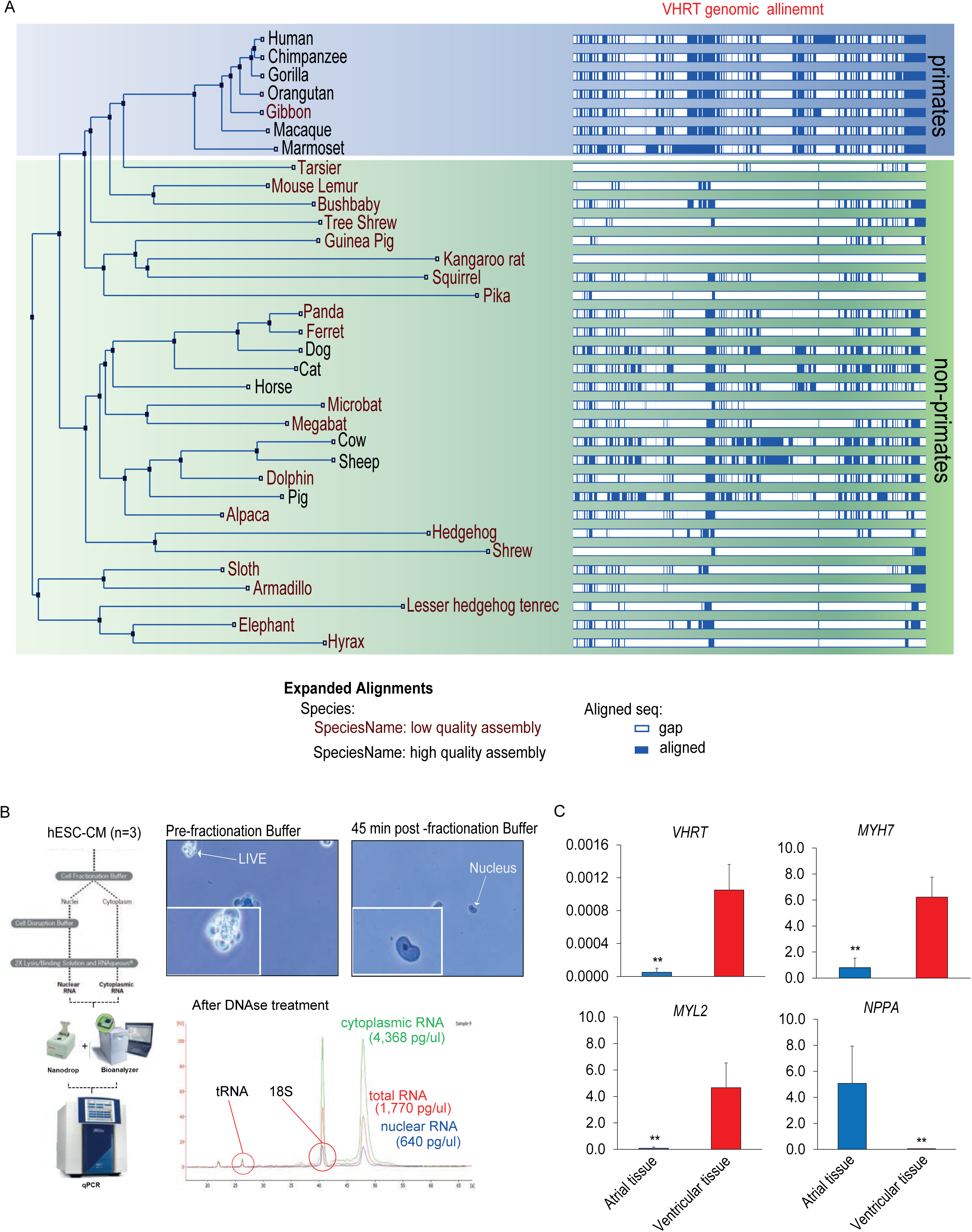
**(A),** Schematic representation showing genomic sequence alignment for *VHRT* extracted from Ensembl Genome Browser^30^. *VHRT* is primarily conserved among the primates. **(B),** Schematic representing the workflow for cellular fractionation. Fractionation efficiency was visualized using trypan blue staining. RNA was extracted from each fraction and treated with DNAse and profiled using Bioanalyzer assay. **(C),** Histogram showing qPCR results for human atrial and ventricular heart tissue for *VHRT*, *MYH7, MYL2* and *NPPA*. n=3. ** p ≤ 0.01.

**Supplementary Figure 6.**
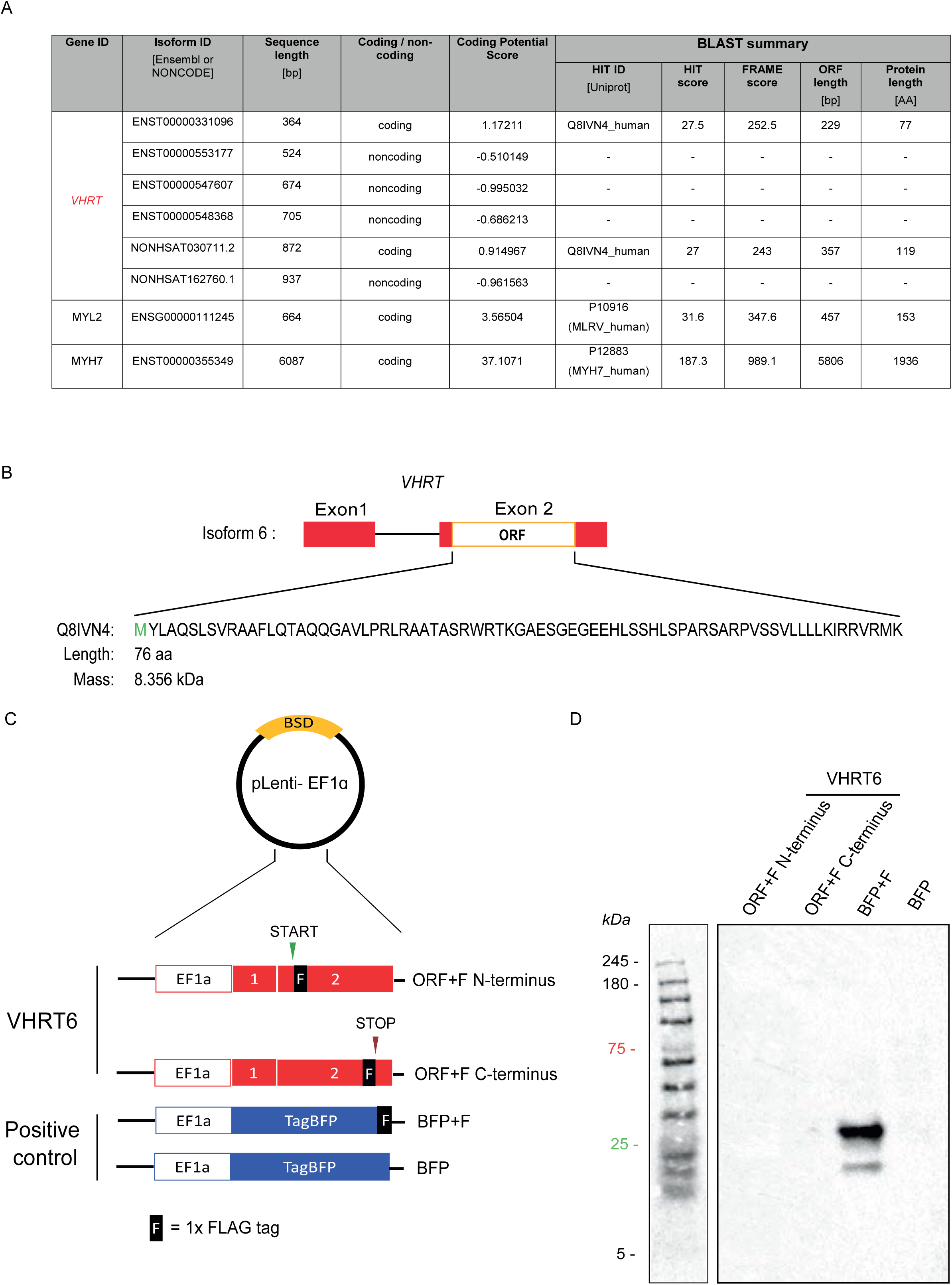
**(A),** Table reflecting the coding potential for each *VHRT* isoform and positive controls using known protein coding genes *MYL2* and *MYH7*. Values were generated using the publicly available coding potential calculator (CPC). HIT ID shows the Uniprot ID related to the gene. A higher HIT score indicates higher likely for protein coding. For a true protein-coding isoform, most hits likely reside within one frame, whereas for a true noncoding isoform, even if it matches certain known protein sequence segments by chance, these random hits are likely to scatter between any of the three reading frames. Thus, FRAME score measures the distribution of the HSPs among three reading frames, the higher the FRAME score, the more concentrated the hits are and the more likely the transcript is protein-coding. **(B),** Schematic showing the small open reading frame (sORF) region on *VHRT6* (*VHRT* isoform 6) and the sequence of the putative peptide with a predicted length of 76 amino acids (aa) and suggested mass of ca. 8 kDa. **(C),** Cartoon showing plasmid and inserts used for *in-vitro* translation. *VHRT6* (without introns) insert was generated with FLAG-tag at the N-terminus or C-terminus of the sORF (in red). Positive control plasmid contained TagBFP (Blue Fluorescent Protein) with FLAG (in blue). All inserts were cloned into lentivirus plasmid (pLenti-EF1α) and used for cell transfection. **(D),** Western blotting using anti-FLAG antibody detecting only a band of ∼ 28kDa for TagBFP positive control plasmid. No bands were detected for *VHRT6*. BFP: blue florescent protein, BSD: blasticidin selection cassette, F: 1xFLAG-tag.

**Supplementary Figure 7.**
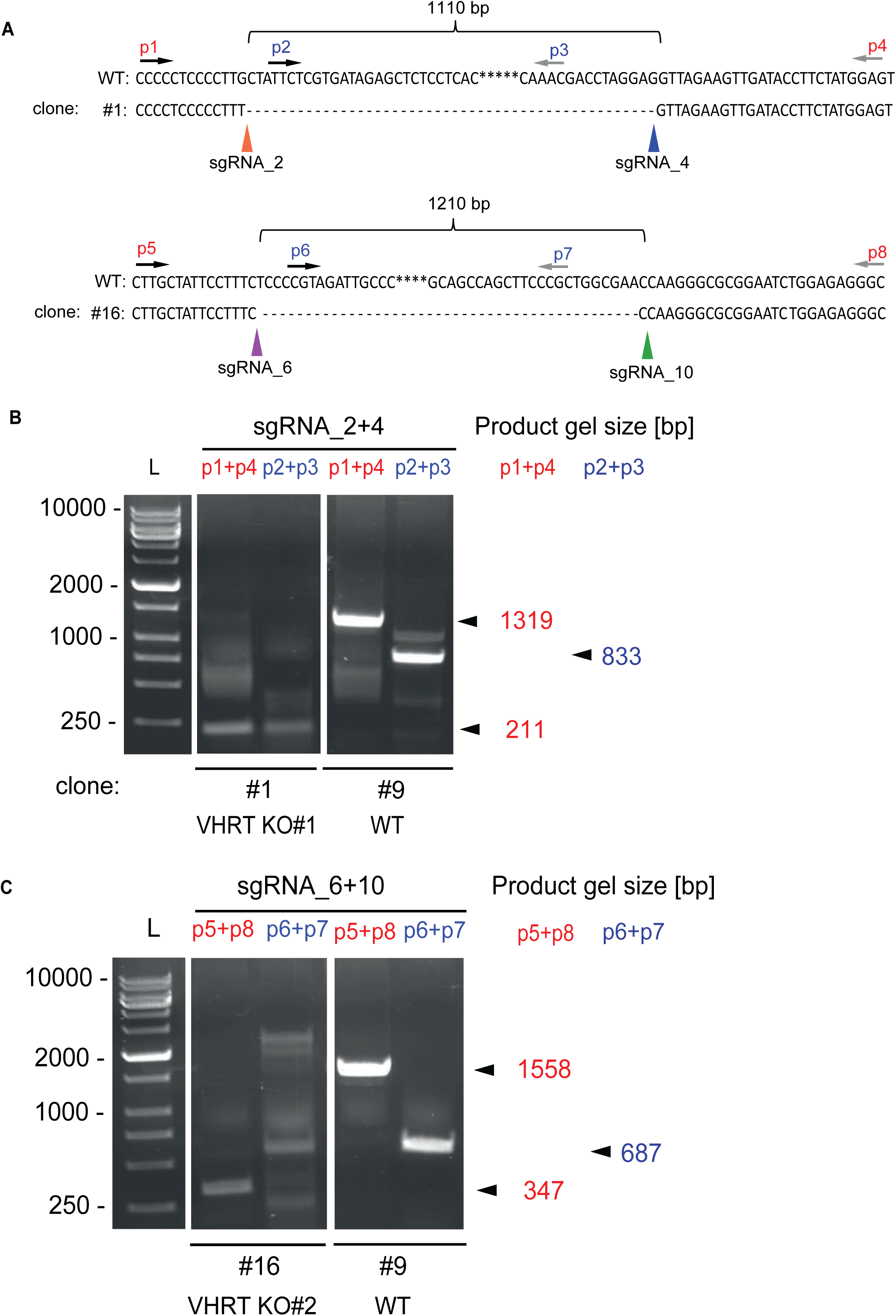
**(A),** Sequence alignment for the deletion of 1110 bp with sgRNA_2/4 set, and 1210 bp with sgRNA_6/10 set. Primers sets used for genotyping are schematically represented. **(B),** Genotyping results proving successful deletion of the *VHRT* locus using sgRNA_2/4 set. Validation primer set p1+p4 and p2+p3, were used and clone #1 resulted in complete *VHRT* KO. **(C),** Genotyping results proving successful deletion of *VHRT* locus using sgRNA_6/10 set. Validation primer set p5+p8 and p6+p7 were used and clone #16 resulted in complete *VHRT* KO.

**Supplementary Figure 8.**
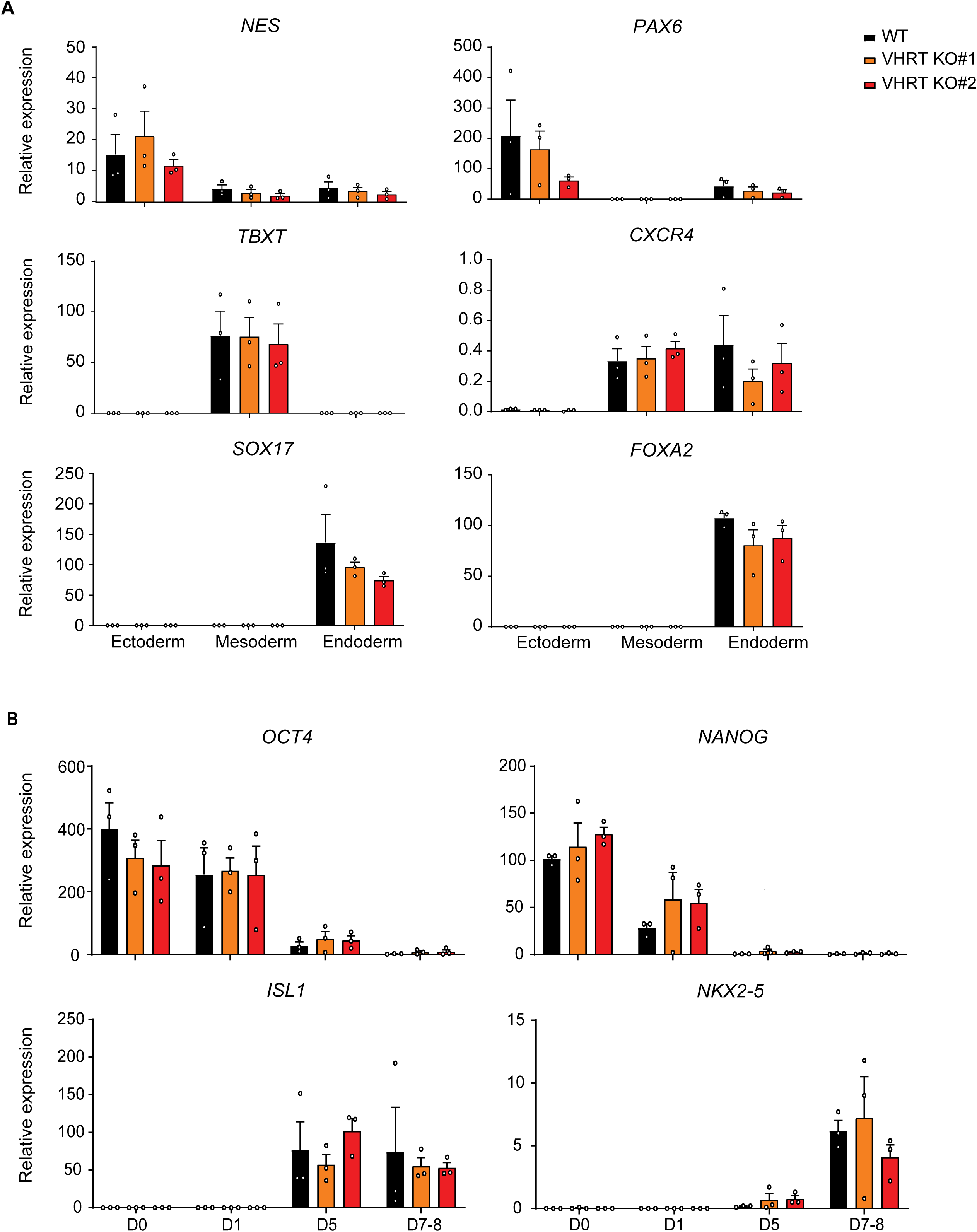
**(A),** Graphs showing the absence of changes in expression of key differentiation markers (*OCT4*, *NANOG*, *ISL1* and *NKX2-5*) in *VHRT* KO cells. **(B),** Tri-lineage differentiation following *VHRT* deletion. Graphs showing absence of significant changes in expression of specific lineage markers, confirming that *VHRT* KO cells were able to successfully specify into ectodermal (*NES* and *PAX6*), mesodermal (*TBXT* and *CXCR4*) and endodermal (*SOX17* and *FOXA2*) lineages despite *VHRT* deletion. Data are represented as fold change expression ± s.e.m, n=3 biological replicates. Student’s paired t-test with a two-tailed distribution.

**Supplementary Figure 9.**
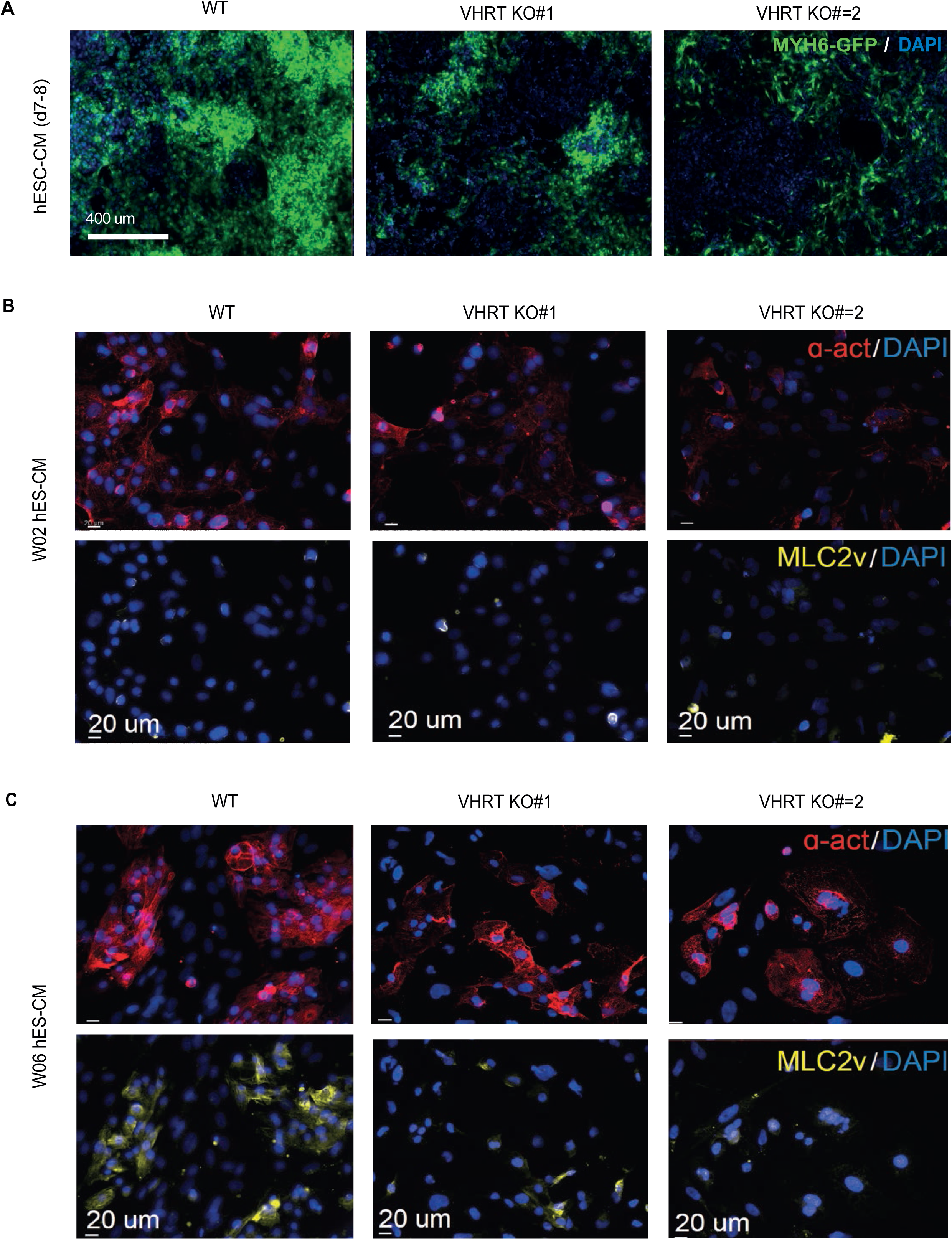
**(A),** Representative images of GFP expression signal at d7-8 post hES differentiation in WT, and *VHRT* KO #1 and #2. **(B),** Immunostaining shows sarcomere disorganization for *VHRT* KO #1 and #2, compared to WT, following W02 and W06 of the CM differentiation protocol. ɑ-act: alpha-actinin 2, and MLC2v: Myosin regulatory light chain 2.

**Supplementary Figure 10.**
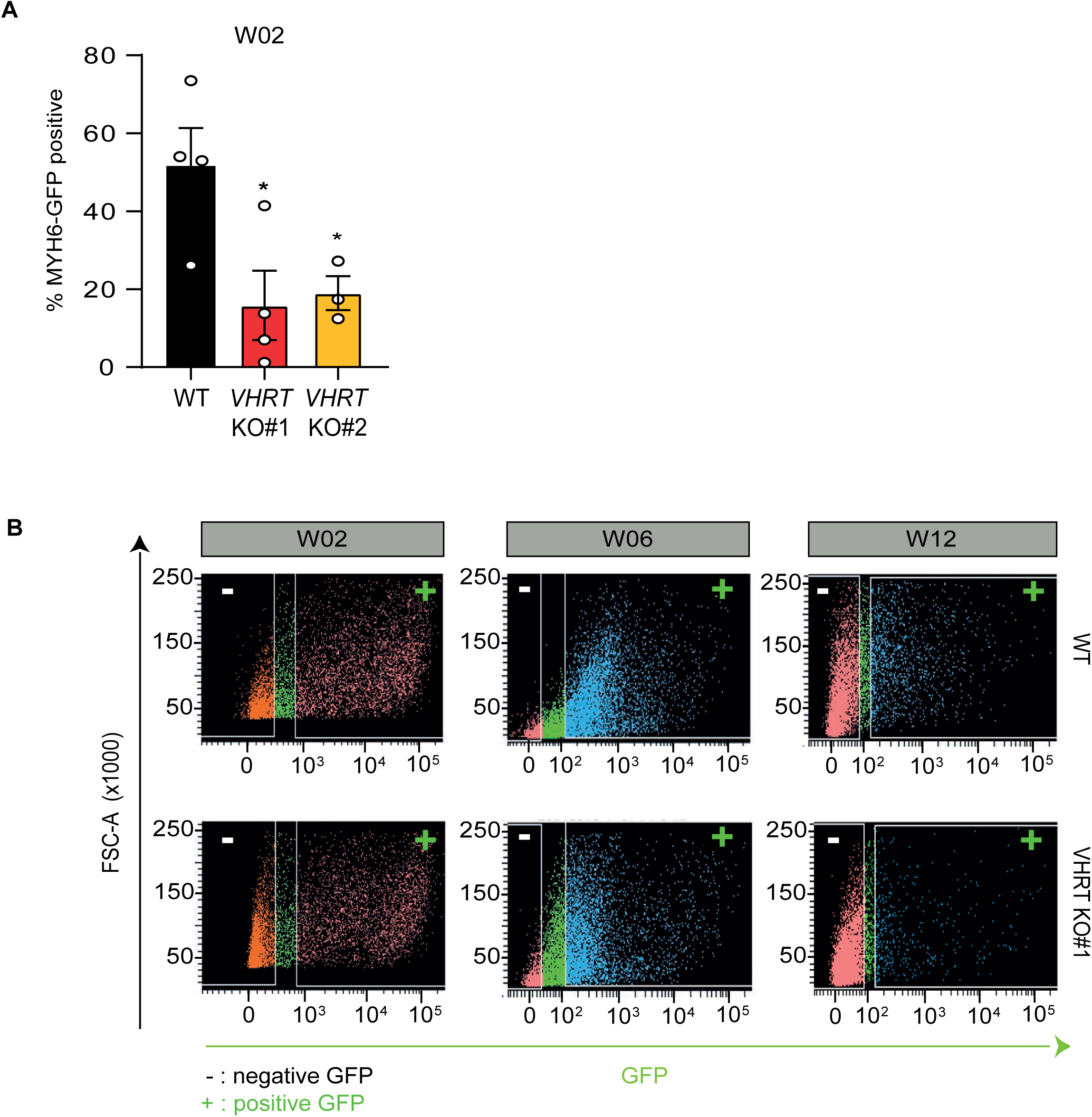
**(A),** Histogram showing the differentiation efficiency as *MYH6*-GFP %positive cells between WT and *VHRT* KO at W02. Error bars represented ± s.d. (n=4 for WT, n=4 *VHRT* KO #1, n=3 *VHRT* KO #2). **(B)** Assay for *MYH6*-GFP positive (+) cells in WT and *VHRT* KO #1 by flow cytometry at W02, W06 and W12.

**Supplementary Figure 11.**
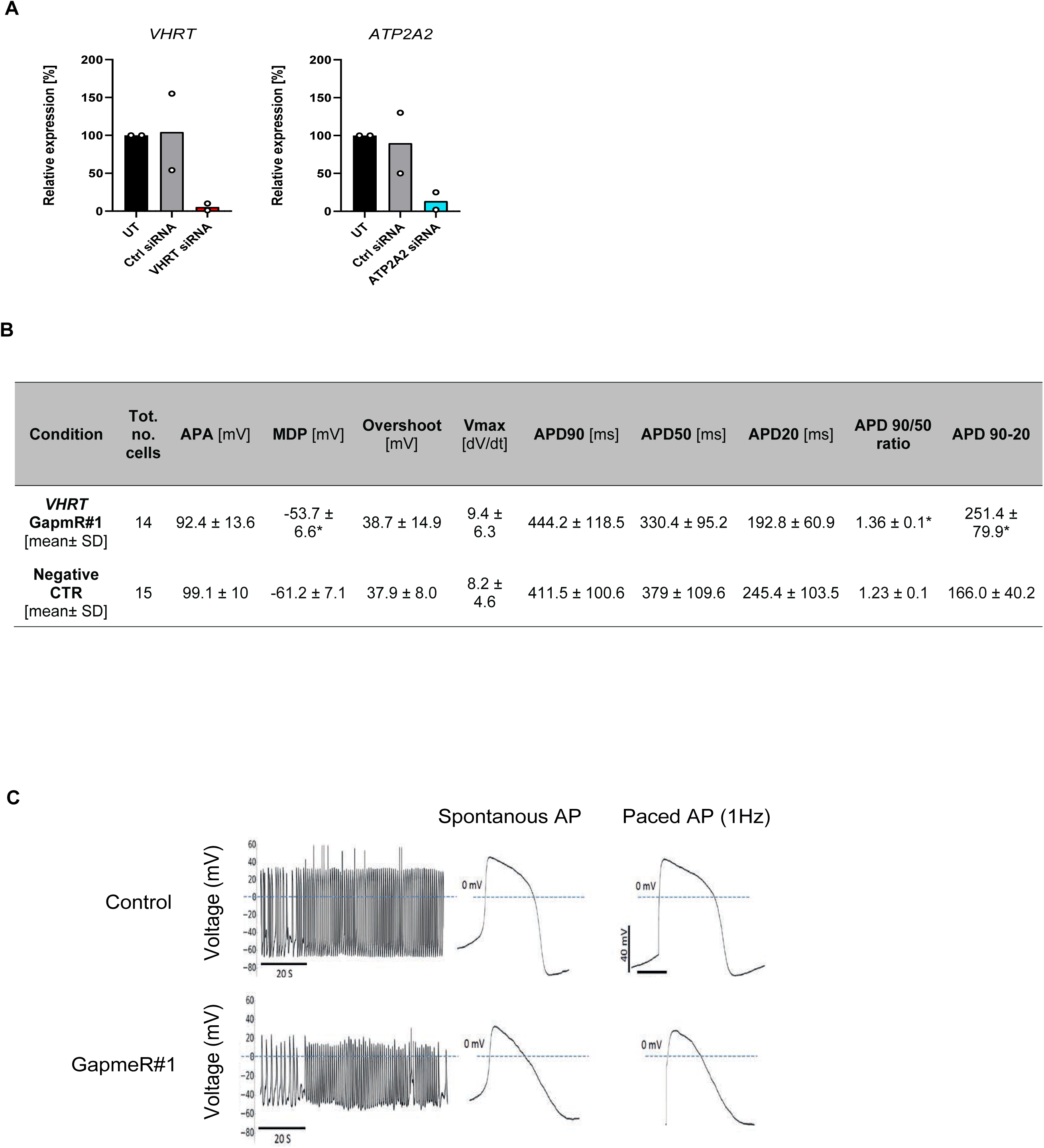
**(A),** Histogram showing qPCR results for *VHRT* and *ATP2A2* siRNA KD (n=2). **(B),** Table showing action potentials recordings. 14 cells transfected with *VHRT* GapmeR#1(*VHRT* knockdown) and 15 cells transfected with Negative A GapmeR (Negative Ctrl) were recorded. hESC-CMs were co-transfected with BLOCK-iT™ Alexa Fluor ® Red Fluorescent to identified positively transfected hESC-CMs. Recordings were made from spontaneously beating cells and with electrical stimulation at 1 Hz. After *VHRT* GapmR#1 knockdown, hESC-CMs showed a less negative maximal diastolic potential (MDP) and a significantly higher APD90/APD50 ratio and APD90-APD20 than Negative Ctrl, indicating an action potential that was less ventricular in nature, compared to negative control cells. Parameters of APs include: AP amplitude (APA), maximal diastolic potential (MDP), overshoot, upstroke velocity of depolarization (Vmax), AP durations (APD) at 20%, 50% and 90% of repolarization (APD20, APD50, and APD90). * p<0.05 compared to Negative A control cells. Student’s t test. **(C),** Representative action potential profile from W06 hESC-CMs after *VHRT* GapmeR#1 KD (lower panel) compared to CTR (non-targeting GapmeR) cells (lower panel).

**Supplementary Figure 12.**
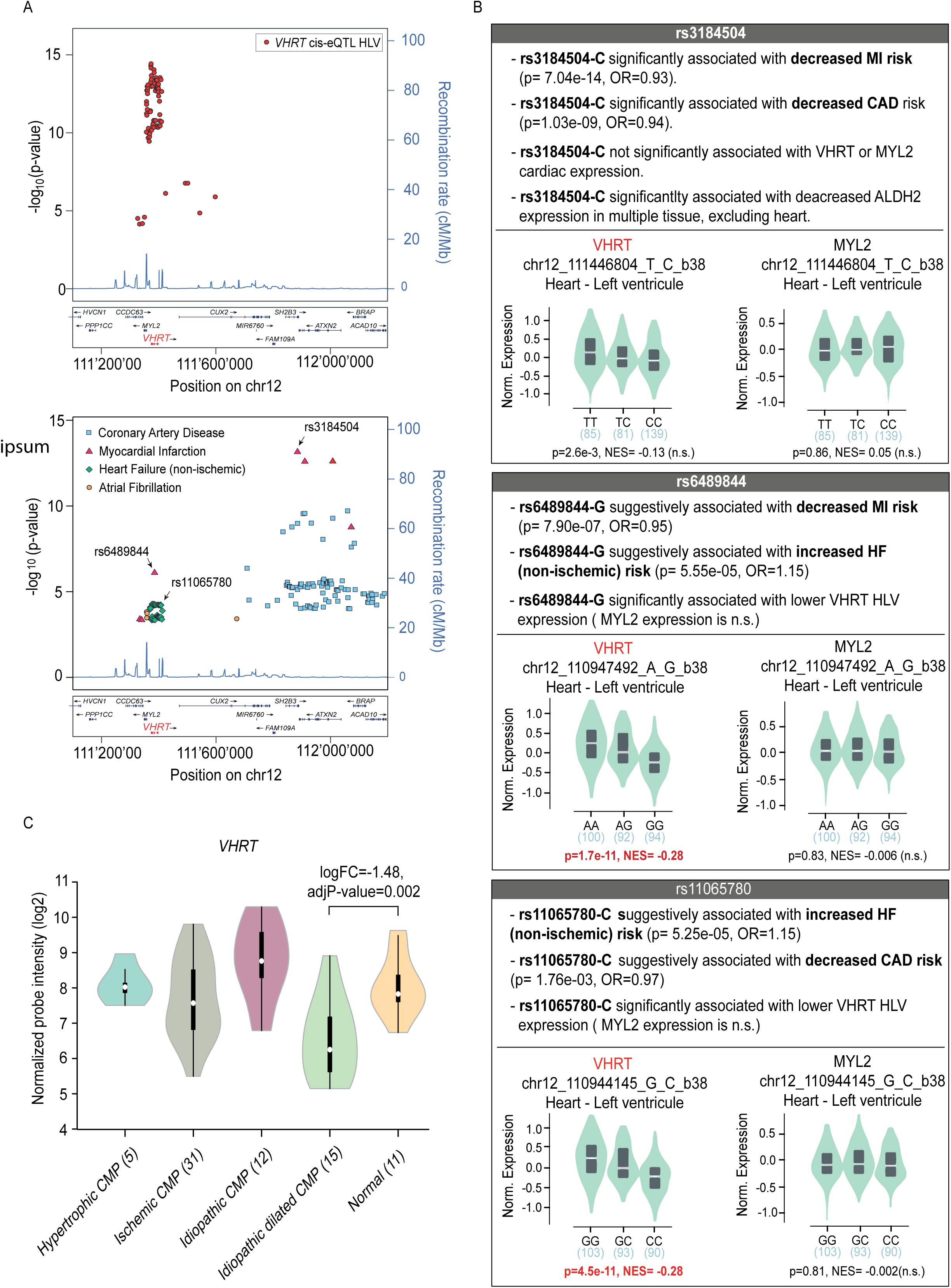
**(A)**, LocusZoom of the 1-MB genomic region harboring *VHRT* depicting significant *VHRT* cis-eQTL SNPs in heart left ventricle (HLV) (upper panel), as well SNPs significantly and/or suggestively associated with CAD, MI, HF (non-ischemic) and/or AF (lower panel). The latter data is taken from published GWAS^8, 9, 11, 19, 20^. **(B),** Top panel showing an example of eQTL rs3184504-C associated with decreased MI and CAD risk, but not with *VHRT* expression. Mid panel displaying the eQTL rs6489844-G, associated with decreased expression of *VHRT* in HLV tissue and suggestively associated with increased risk for HF, but decreased risk of MI. Similarly, on the bottom panel the eQTL rs11065780-C is significantly associated with low *VHRT* expression in HLV and increased HF, but suggestively associated with decreased CAD risk (bottom panel). NES: normalized effect size. P < 0.05. **(C)**, *VHRT* (probeset 230195_at) is significantly downregulated in left ventricular (LV) tissue in patients with Idiopathic Dilated Cardiomyopathy (N=15) versus healthy controls (N=11), but not in Hypertrophic-(N=5), Idiopathic (N=12) or Ischemic (N=31) cardiomyopathy based on reanalysis of microarray data from GEO study GSE1145 (https://www.ncbi.nlm.nih.gov/geo/query/acc.cgi?acc=GSE1145).

**Supplementary Figure 13.**
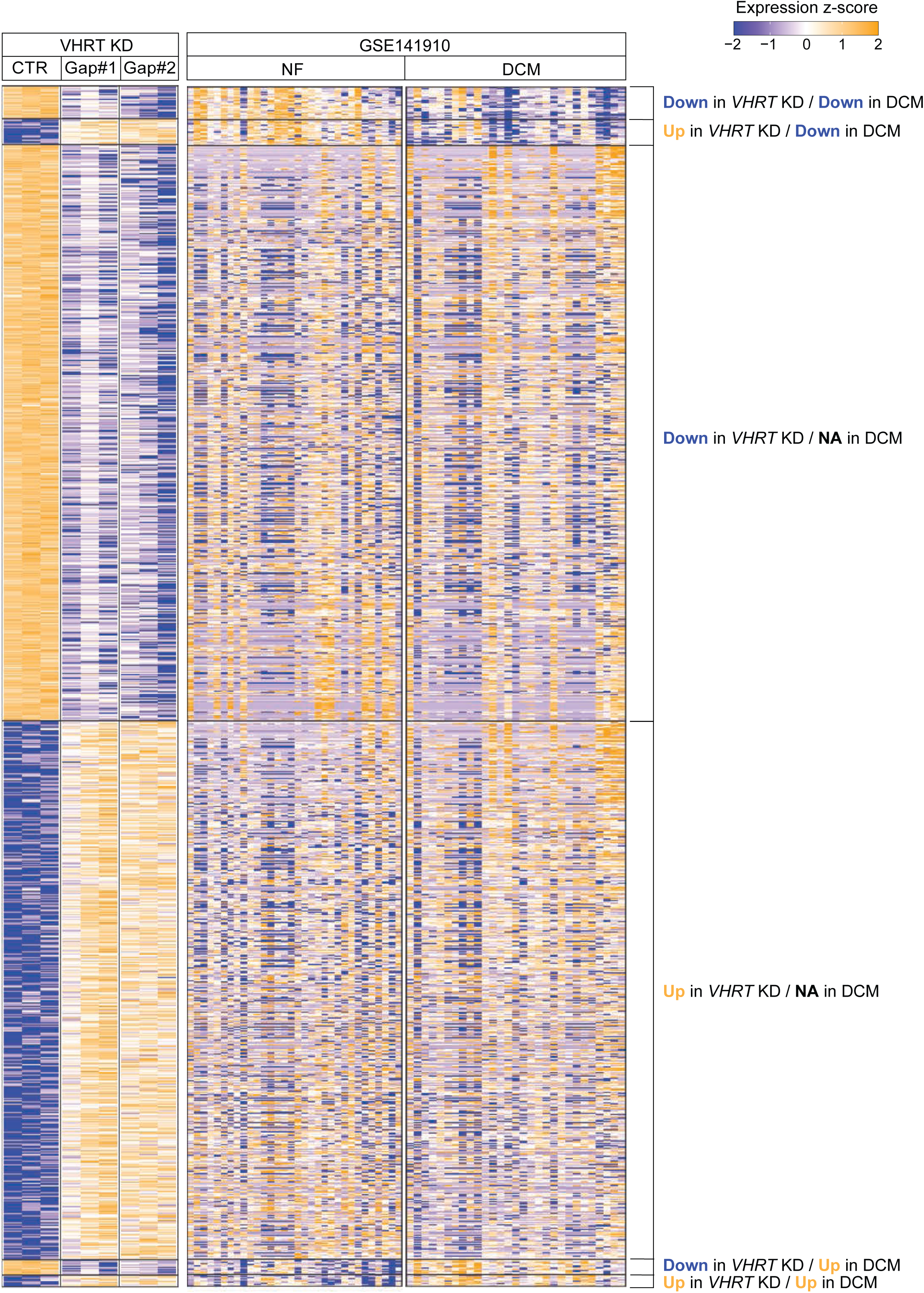
**A)**, Heatmap of normalized expression values of 1100 significantly differentially expressed (DE) genes in *VHRT* KD, compared to the RNAseq from Non-Failing (NF) versus failing (Dilated Cardiomyopathy, DCM) hearts. The subset that were consistent for downregulation in the 2 comparisons are represented in Figure 6.

## SUPPLEMENTARY TABLES

**Supplementary Table 1:** Single cell RNAseq and data quality.

**Supplementary Table 2:** Tabulation of gene expression in each week.

**Supplementary Table 3:** WGCNA module and gene kME values.

**Supplementary Table 4:** Gene ontology of 6 modules.

**Supplementary Table 5:** VHRT transcripts description and annotation.

**Supplementary Table 6:** Gene ontology for genes differentially regulated in VHRT WT vs VHRT KO #1 in 02W,06W,12W old hES-CM.

**Supplementary Table 7:** Gene ontology for genes differentially regualted in VHRT CTRL vs VHRT GapmeR KD.

**Supplementary Table 8:** GeneNetwork data analysis.

**Supplementary Table 9:** Human Dilated cardiomyopathy (DCM) and non-failing (NF) tissue biodata.

**Supplementary Table 10:** Combined transcriptome analysis of VHRT GapmeR KD and human Dilated cardiomyopathy.

**Supplementary Table 11:** Oligonucleotides.

